# Nucleocapsid mutation R203K/G204R increases the infectivity, fitness and virulence of SARS-CoV-2

**DOI:** 10.1101/2021.05.24.445386

**Authors:** Haibo Wu, Na Xing, Kaiwen Meng, Beibei Fu, Weiwei Xue, Pan Dong, Yang Xiao, Gexin Liu, Haitao Luo, Wenzhuang Zhu, Xiaoyuan Lin, Geng Meng, Zhenglin Zhu

## Abstract

In addition to the mutations on the spike protein (S), co-occurring mutations on nucleocapsid (N) protein are also emerging in SARS-CoV-2 world widely. Mutations R203K/G204R on N, carried by high transmissibility SARS-CoV-2 lineages including B.1.1.7 and P.1, has a rapid spread in the pandemic during the past year. In this study, we performed comprehensive population genomic analyses and virology experiment concerning on the evolution, causation and virology consequence of R203K/G204R mutations. The global incidence frequency (IF) of 203K/204R has rose up from nearly zero to 76% to date with a shrinking from August to November in 2020 but bounced later. Our results show that the emergence of B.1.1.7 is associated with the second growth of R203K/G204R mutants. We identified positive selection evidences that support the adaptiveness of 203K/204R variants. The R203K/G204R mutant virus was created and compared with the native virus. The virus competition experiments show that 203K/204R variants possess a replication advantage over the preceding R203/G204 variants, possibly in relation to the ribonucleocapsid (RNP) assemble during the virus replication. Moreover, the 203K/204R virus increased the infectivity in a human lung cell line and induced an enhanced damage to blood vessel of infected hamsters’ lungs. In consistence, we observed a positive association between the increased severity of COVID-19 and the IF of 203K/204R from in silicon analysis of global clinical and epidemic data. In combination with the informatics and virology experiment, our work suggested the contribution of 203K/204R to the increased transmission and virulence of the SARS-CoV-2. In addition to mutations on the S protein, the mutations on the N protein are also important to virus spread during the pandemic.

## Introduction

Since the outbreak of the severe acute respiratory syndrome coronavirus 2 (SARS-CoV-2) globally from 2019, COVID-19 has caused more than 160 million confirmed infections and more than 3 million deaths worldwide. SARS-CoV-2 is rapidly evolving RNA virus and causes severe pneumonia. The virus threat the life of older age groups and those with chronic conditions. Due to the genetic proofreading mechanism ^1^ and error prone replication, SARS-CoV-2 ^2^ does not have a high mutilation rate ^3^. However, short generation time and large population size enabled this virus to evolve rapidly, and mutates continually during transmission. These genomic variations may contribute to the severity of disease and the efficiency of transmission.

In general, most viral mutations are deleterious to the virus and disappear soon, whereas mutations that persist and increase in frequency may be selectively neutral or advantageous to viral fitness. However it is challenging to classify the mutation to neutrality or positive selection. Particularly for a newly emerging virus such as SARS-CoV-2, a new increasing mutation may be a result of neutral epidemiological processes such as genetic bottlenecks following founder events and range expansion. Among all the protein expressed by the virus, the spike (S) protein directly interacts with the humane ACE2 protein ^4^ and is the target for vaccines and therapeutics ^5^. Thus previous research has been focused on adaptive SARS-CoV-2 mutants on the spike glycoprotein (S), such as D614G, N501Y ^6^ and E484K ^7^. The D614G mutants have a dramatic increase in incidence frequency (IF) and an increased fitness, infectivity and fatality ^8–12^. The mutant spread rapidly in the first half year of 2020 and became predominant in population after Jul-2020 ^13^. N501Y was identified to be capable of increasing the virus infectivity by 52% ^14^. Moreover, 501Y showed high resistance to neutralization ^15^, indicating the escape of neutralization by vaccine. The lineage with two or more adaptive mutants may have higher fitness than the one with only one adaptive mutant.

The growing lineage B.1.1.7^16^ carrying N501Y (N501Y.v1) is identified to be with increased infectivity and capability to reduce the activity of neutralization ^17^. Following B.1.1.7, another lineage B.1.351 ^18^ was identified and considered the second version of N501Y variants (N501Y.v2). This lineage includes another S mutation E484K, which is capable of evading the neutralization of most monoclonal antibodies ^19^. The lineage P.1 ^20^, also includes 484K, is the third version of N510Y (N501Y.v3). P.1 and B.1.315 are lineages resistant to antibodies ^21^. B.1.429+B.1.427 and B.1.525 are two newly identified lineages also associated with immune evasion ^22^. These rapidly growing lineages all contain D614G, which is associated with the enhanced transmission ^23^. Except for D614G, N501Y ^6^ and E484K ^7^, there are other mutants that distinguish these lineages from the original virus. The evaluation of these mutants for functional effects is important.

R203K/G204R is a co-occurring N mutation, growing rapidly and with potential association with the infectivity of virus ^10^. This mutation carried by B.1.1.7 and P.1 referred above was identified with selection signatures by our previous work ^10,24^. We kept tracking the evolution of R203K/G204R based on all documented SARS-CoV-2 genome sequences monthly, and found this mutation has a second time rapid growth accompanied with the emergence of B.1.1.7 (Figure 1). The mutation R203K/G204R are becoming dominant in the worldwide pandemic, and may have positive effects to the fitness of SARS-CoV-2. Thus, a thorough evaluation of the evolutionary and functional effects of R203K/G204R is important for our understanding the effects of N mutations and the contribution of N mutations to present rapidly growing lineages. We constructed the R203K/G204R mutant virus by single point mutation. Through experimental investigation in cell lines, hamsters and a human airway tissue model, we identified and validated an increased infectivity and fitness of 203K/204R variants. In hamster lung tissues, we observed an increased severity caused by 203K/204R virus, confirmed by epidemic surveys and statistics of global clinical data. By experiments and structural prediction, we concluded that it is possibly the change of the N protein charge that enforced an enhanced virus replication, which eventually resulted in the increase of infectivity and fitness. These indicated that the N mutation R203K/G204R should be associated with the increased transmission ^25^ and virulence ^26^ of B.1.1.7 and P.1. R203K/G204R deserves more attention in the future.

**Figure 1.**
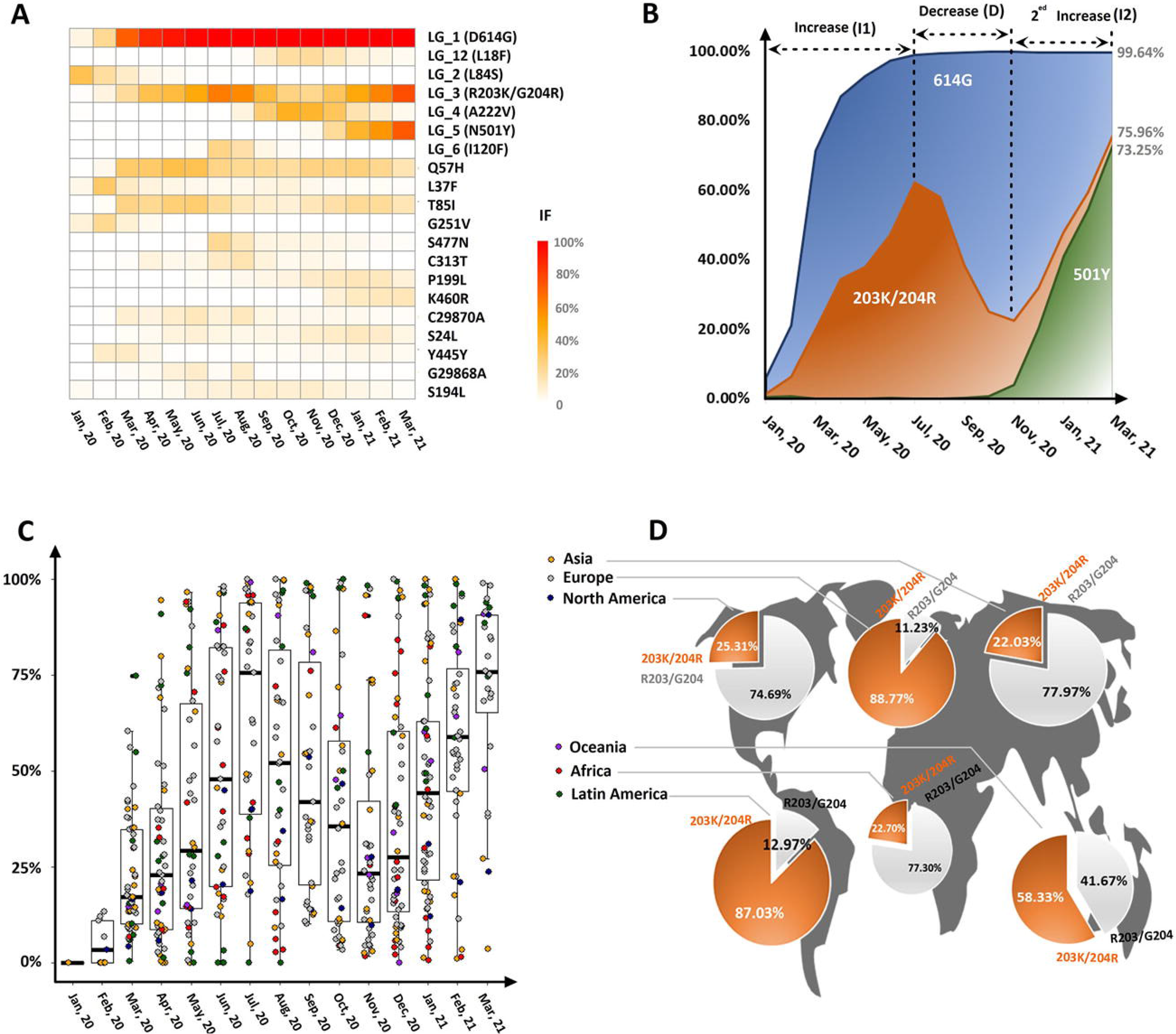
Rapid spread of R203K/G204R. (A) Heatmap showing the global IF changes by continuous colors from white (0%) to red (100%). Y are LGs/mutations with an IF > 10% in one month. (B) Global IF tracks of the top three mutants (614G, 510Y and 203K/204R) with the highest IFs to date. The time intervals, when the IF of 203K/204R increased (I1), decreased (D) and increased for the second time (I2), are annotated at the top. (C) IF change in countries over months. The continents of countries are differentiated by colors. (D) IFs of 203K/204R (orange) and R203K/G204 (gray) in continents.

## Result

### The rapid spread of R203K/G204R worldwide

We performed population genomic analyses of 884736 full-length SARS-CoV-2 genomes (Table S1) collected from GISAID ^27^ to track the changes of SARS-CoV-2 mutations. We found 96 mutations with a monthly incidence frequency (IF) higher than 0.05 (Figure S1). Through pairwise linkage disequilibrium analyses of these 96 mutation sites (Table S2, S3, Figure S2), we identified 12 mutation linkage groups (LGs), named LG_1 to LG_12 in the following description for convenience. There are 26 singleton mutations, without an association with any LG. LG_3 has three adjacent nucleotide mutations, from 28881 to 28883, completely linked (ρ^2^ > 0.99, Table S2). The corresponding amino acid change in protein level is a co-occurring R203K/G204R in the nucleocapsid (N) protein. We found R203K/G204R has a rapid spread worldwide. Its global IF has increased from nearly zero in Jan-2020 to more than 70% in Mar-2021 (Figure 1A, B). We tracked the IFs of 203K/204R in countries (Figure 1C) and observe a consistent IF track (compared to Figure 1B, correlation = 0.96, P-value = 1.41e-08). Up to Mar-2021, 203K/204R has token a percentage higher than 80% in Europe and Latin America (Figure 1D). In intra-host single nucleotide variations (iSNVs) analyses, we also observed a significant IF increase of R203K/G204R (Figure S3).

### The competition and cooperation of 203K/204R and other mutants

In the first half year of 2020, there is a fast increase of IF both for 614G (LG_1) and 203K/204R (LG_3) (Figure 1B). To identify if the IF increase of 203K/204R is resulted from purifying selection led by 614G (LG_1), we plotted the IF tracks of four combinations, D614/R203, D614/203K, 614G/R203 and 614G/203K. We found 203K has an IF increase not only in D614 but also in 614G virus (Figure S4). The IF track of 614G/203K (Figure S4B) has a high correlation (1, P-value < 2.2e-16) with the IF track of 203K/204R (Figure 1B). There is no linkage disequilibrium between D614G and R203K/G204R (ρ ^2^ = 0.043, Table S2). These indicated that the evolution of R203K/G204R is independent from D614G.

The IF change of 203K/204R is not in one direction all the time. It encounters an increase (I1) from Jan-2020 to Jul-2020, a decrease (D) from Jul-2020 to Nov-2020, and an increase for the second time (I2) from Nov-2020 to date (Figure 1B). For the causation of the decrease, we performed correlation tests of IF tracks between pairs of mutants in the three time intervals, I1, D and I2 (Figure 1B). R203K/G204R has a significant positive correlation (r+) with C313T in I1, a significant negative correlation (r-) with LG_4 and an r+ with LG_6 in D. In I2, R203K/G204R has an r- with LG_4, an r- with S194L, an r+ with LG_5 and an r+ with S477N (Figure S5). We counted the frequencies of those six LGs/mutations in R203/G204 variants and in 203K/204R variants (Figure S6A, B). We also counted the frequencies of R203/G204 and 203K/204R in the six LGs/mutants (Figure S6C-H). We observed a significant correlation between the IF track of 222V (LG_4) in R203/G204 virus and the IF track of R203/G204, at the time intervals D and I2 (correlation = 0.96, P-value = 2.933e-05, Figure S6A). R203/G204 is predominant in 222V (LG_4) (Figure S4C). The rise of 222V (LG_4) in R203/G204 variants may lead to the decrease of 203K/204R.

There is a sharp rise of 501Y (LG_5) in 203K/204R variants at I2. There is also a significant correlation between the IF change of 501Y (LG_5) in 203K/204R virus and the IF change of 203K/204R at I2 (correlation = 0.99, P-value = 0.00016, Figure S6B). 501Y (LG_5) is always low in percentage in R203/G204 virus (Figure S6A).

At most time, 203K/204R is predominant in 501Y (LG_5) mutants (Figure S6D). The co-occurring of 501Y and other mutants in LG_5 and 203K/204R may lead to the second IF increase 203K/204R. 120F, 313T and 477N possibly emerged in 203K/204R strains and grew at I1, but shrank thereafter along with the IF decrease of 203K/204R (Figure S6A, B, E-G). The combination of these mutants and 203K/204R may have lower fitness than the combination of 222V and R203/G204. 120F, 313T, 477N and 194L have low IFs (<30%) over time. Their effects to 203K/204R is ignorable.

There are in total 2^3^ = 8 possible combinations for three two-allele polymorphisms.

For a further understanding the effects of A222V (LG_4) and N501Y (LG_5) to 203K/204R, we tracked the IFs of those 8 combinations, in which we identified four dominant lineages (Figure 2A). R203/G204 is carried by A222 + N501 + R203/G204 (ANR) and 222V + N501 + R203/G204 (VNR), while 203K/204R is carried by A222 + N501 + 203K/204R (ANK) and A222 + 501Y + 203K/204R (AYK). The IFs of lineages (Figure 2A) show that the order of adaptiveness from high to low possibly is AYK, VNR, ANK and ANR. For the four lineages, the global IF change is in consistent with the medians of IFs in countries (Correlations > 0.89, P-value < 0.05, Figure S7). The rise of VNR mostly happened in European countries (Figure S8). In UK (Figure 2B), the rise of VNR is accompanied by the shrinking of ANR and ANK. Later, with the rise of AYK, VNR, ANR and ANK vanished (Figure 2B). In Mexico and India (Figure 2C, D), there is a continuous growing of ANK and shrinking of ANR. ANK and AYK has a slower spread in North America countries, e.g. USA (Figure 2E, S8), than in Europe countries, e.g. UK (WilCox Test, P-value = 0.00017). In South Africa, AYR spread rapidly and finally replaced the preceding ANR and ANK (Figure 2F). The AYR variants are mostly B.1.351 (1211/1379, 87.8%).

**Figure 2.**
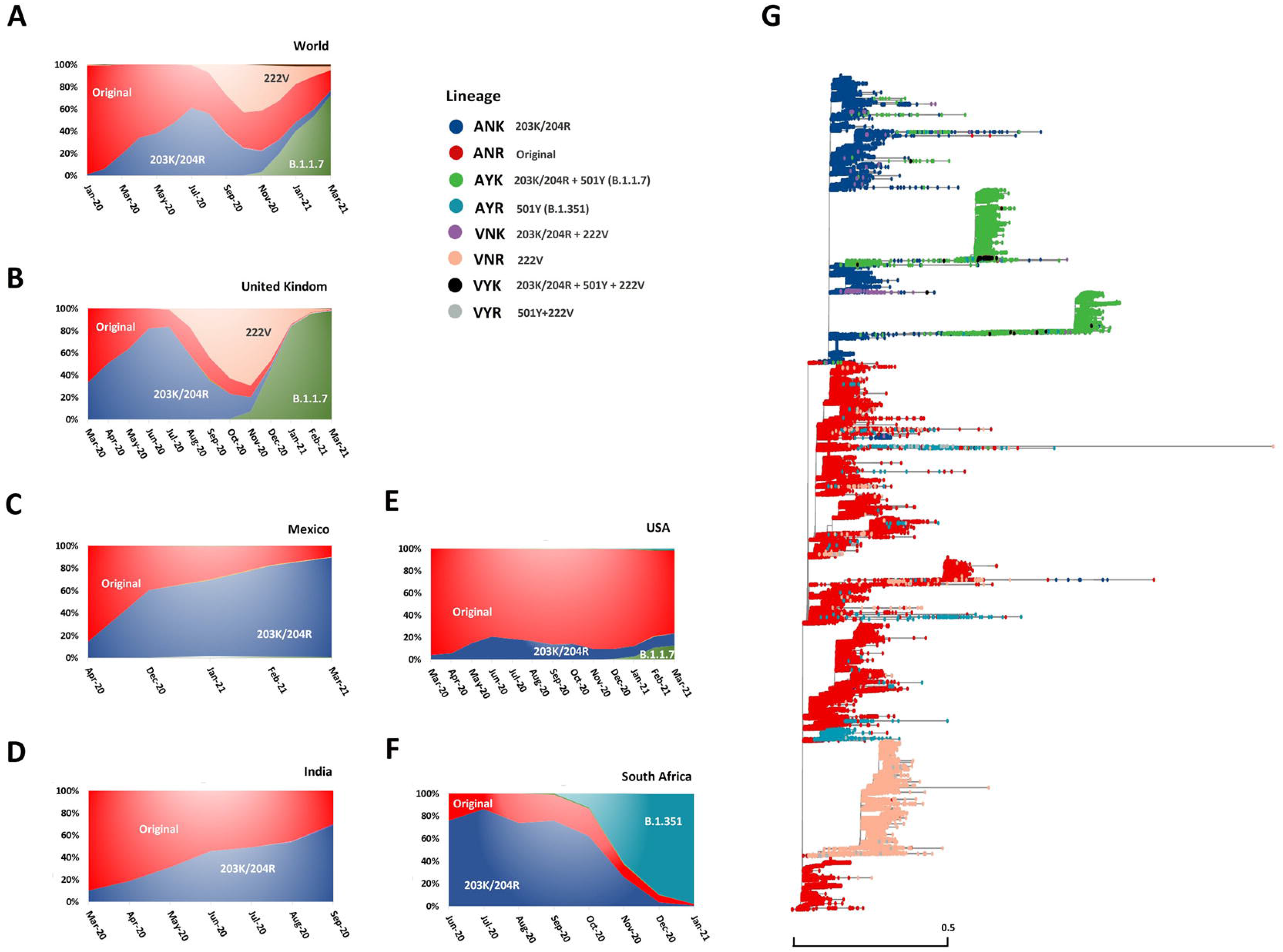
IF changes of 8 lineages covering A222V (LG_4), N510Y (LG_5) and R203K/G204R (LG_3) and the distribution of lineages in the phylogenetic tree of all strains (G). A to F are the percentage (IF) accumulated area map of lineages in the world and five countries, respectively. In B to F, the IFs with insufficiency of sample size (< 200) are not shown. Lineages are differentiated by colors. The legend describing lineages is in the center. The third column of the legend is to highlight the mutants carried by each lineage. ‘Original’ denotes the co-existence of the original allele (A222, N501 and R203/G204).

### The emergence of B.1.1.7 in 203K/204R mutants

In the lineages with a rapid growth (Figure S9, S10), B.1.1.7 and P.1 carry 203K/204R (Table S4). B.1.1.7 has 28 mutations, including 4 LG_1 (D614G) mutations, 3 LG_3 (R203K/G204R) mutations and 21 LG_5 (N501Y) mutations (Table S4). The LG_1 (D614G) variants are fixed and B.1.1.7 is just the combination AYK referred above. The crossing of B.1.1.7 and AYK (216578 strains) take a percentage of 95.4% for all strains (226992) belonging to B.1.1.7 or AYK. Most 501Y evolved in 203K/204R virus (Figure S6B). For a further identification of the relationship between 203K/204R and B.1.1.7, we constructed a phylogenetic tree using all SARS-CoV-2 strains (Table S1). The distribution of lineages along the tree showed that the origin of B.1.1.7 (AYK) is ANK (Figure 2G, S11). The construction of a TSC network of all strains (Figure S9) also showed that AYK evolved from ANK. We created animations to display the evolution of SARS-CoV-2 lineages in countries (Supplementary Animations, the legends follow Figure 2G). These animations indicated that the emergence of AYK in ANK is multiregional.

### Selection signatures for R203K/G204R

The IF increase of 203K/204R infers that the adaptiveness. For evaluation, we performed sliding window calculation of the composite likelihood ratio (CLR) ^28,29^ in SARS-CoV-2 genomes. A CLR peak surpassing the threshold (the top 5% CLR in ranking) is a positive selection signature with statistical significance. For convenience in manipulation, we calculated the ratio (m/t) of the CLR at the mutation site (m) and the CLR threshold (t) in the genome. Thus, a CLR _m/t_ equal or higher than 1 means a significant CLR peak or an adaptive selection signature. We calculated the CLR _m/t_ for R203/G204 variants and 203K/204R variants, respectively. Considering there are pattern changes in diversification ^30^ and the spread of R203K/G204R is not continuous, we calculated the CLR _m/t_ per month. We identified positive selection signatures. Taking the results from UK strains as an example, there are more CLR peaks (Chisq-test P-value=0.098) for 203K/204R variants than R203/G204 variants (Figure 3A, B). In July, the growth of 203K/204R encountered a peak. In the months near this time point, we observed a significant higher 203K/204R CLR _m/t_ in countries than R203/G204 (Figure 3C, S13A). In the comparison of the whole genome diversification between R203/G204 and 203K/204R variants, we found that Tajima’s D, Pi and Theta are lower in 203K/204R than R203/G204 variants (Figure 3D, S13B-D).

**Figure 3.**
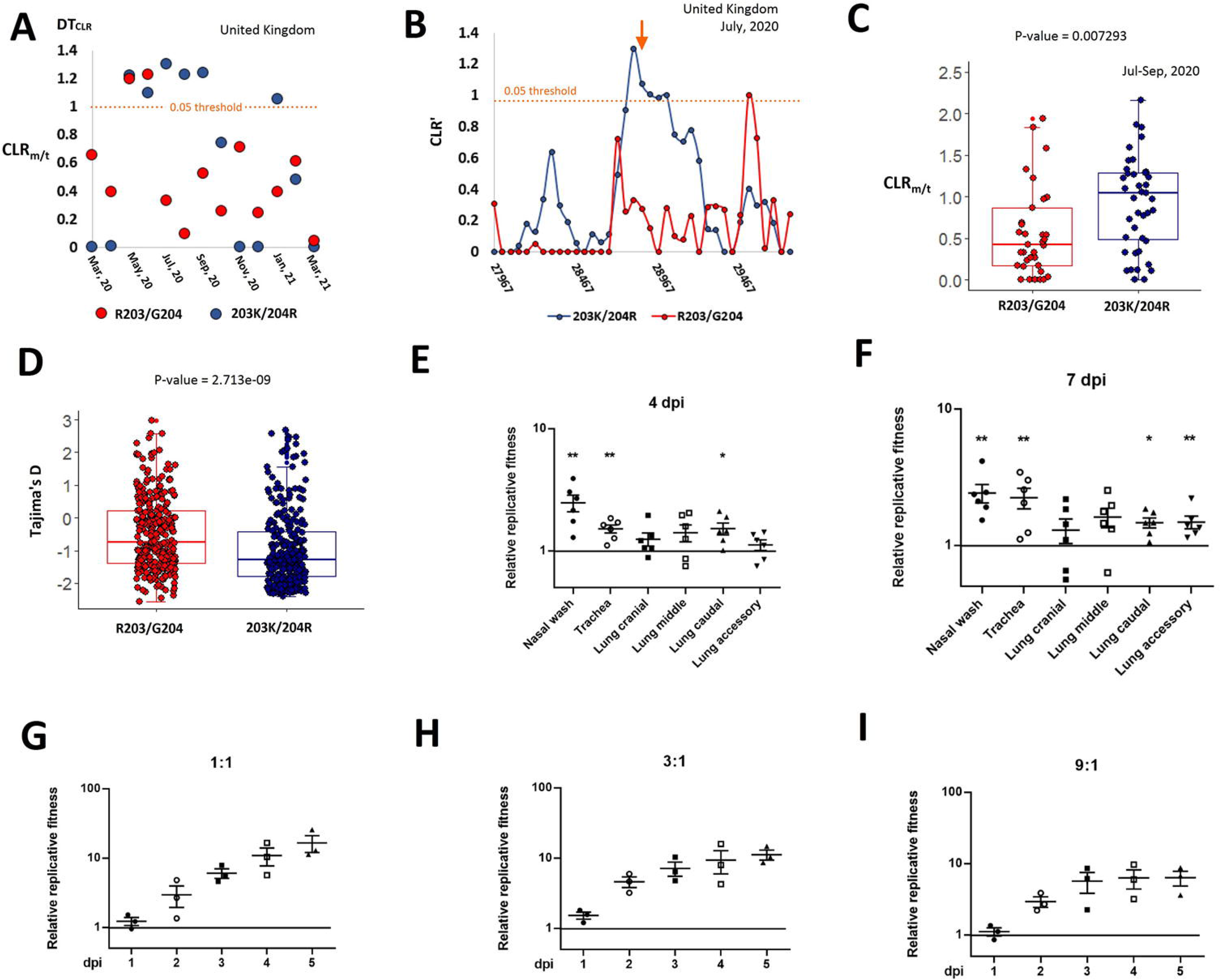
Evidences showing the adaptiveness of R203K/G204R. (A) Comparison of CLR _m/c_ per month in UK between R203/G204 (red) and 203K/204R variants (darkblue). An orange dotted line denotes the top 5% CLR cutoff. (B) A sliding window view showing a CLR peak exceeding the threshold (the orange dotted line). This plot is a detailed view for the positive CLR _m/c_ in July in UK, as shown in (A). CLR’ is a transformed CLR value (CLR’ = CLR / c). The transformation is to diminish the background difference effects. (C) Comparison of CLR _m/c_ in the months from July to September between R203/G204 (red) and 203K/204R variants (darkblue). (D) Comparison of Tajima’s D calculated by months and countries between R203/G204 (red) and 203K/204R viruses (blue). (E, F) Hamsters were inoculated with a 1:1 mixture of R203/G204 and 203K/204R viruses (104 PFU each). Nasal wash, trachea and lungs were collected on day 4 (E) and 7 (F) after infection. Relative amounts of R203/G204 and 203K/204R RNA were assessed by RT-PCR and Sanger sequencing. Y-axis used a log10 scale. Data were represented as mean ± s.e.m.. (E) and (F) show that 203K/204R virus is advantageous over R203/G204 virus in hamsters. (G-I) Competition assay. A mixture of R203/G204 and 203K/204R viruses with initial ratios of 1:1 (G), 3:1 (H) or 9:1 (I) were inoculated onto human airway tissue cultures at a total MOI of 5. Virus ratios after competition were measured by RT-PCR and Sanger sequencing. Y-axis used a log10 scale. All data were represented as mean ± s.e.m.. *, p<0.05, **, p<0.01. Abbreviation: n.s., no significance.

### 203K/204R show a higher fitness than R203/G204

We constructed the 203K/204R mutant virus and tested the virus replication in hamsters using R203/G204 as a control. We made hamsters infected with a 1:1 mixture of R203/G204 and 203K/204R viruses and then assessed the competition of them at 4 days and 7 days after infection. We observed a higher 203K/204R to R203/G204 ratio, indicating a replication advantage for 203K/204R in hamsters (Figure 3E, F). The ratio of 203K/204R to R203/G204 was greater than 1 on day 4 and 7 after infection (dpi), indicating that 203K/204R virus had a consistent advantage over R203/G204 virus (Figure 3E, F).

Moreover, we performed competition experiments in the human airway tissue culture model. After infecting the tissues with a 1:1 infectious ratio of R203/G204 and 203K/204R viruses, the 203K/204R to R203/G204 ratio was increased from 1 dpi to 5 dpi (Figure 3G). In addition, after infecting the airway culture with 3:1 or 9:1 ratio of R203/G204 and 203K/204R viruses, the 203K/204R virus rapidly overcame its initial deficit and reached an advantage over R203/G204 virus (Figure 3H, I). These results demonstrated that the 203K/204R virus could rapidly outcompete the R203/G204 virus when infecting human airway tissues.

### 203K/204R has a higher infectivity than R203/G204

We compared the infectivity of the constructed mutant and the wild virus in cell lines. We found the 203K/204R virus replicated with a higher infectious titre than R203/G204 at 36 h post-infection (hpi) in Vero E6 cells (Figure 4A). A similar trend of extracellular viral RNA production was observed in infected Vero E6 cells, too (Figure 4B). To compare the infectivity, we calculated the genomic RNA/plaque-forming units (PFU) ratio. We found no significant differences in virion infectivity between the two viruses in Vero E6 cells (Figure 4C). Next, we compared the replication kinetics of R203/G204 and 203K/204R viruses in human lung epithelial cell line Calu-3. With a multiplicity of infection (MOI) of 0.01, 203K/204R produced more infectious virus at 36 and 48 hpi than R203/G204 (Figure 4D), indicating that 203K/204R enhanced viral replication. However, the cells infected with 203K/204R virus produced almost equivalent extracellular viral RNA compared to cells infected with R203/G204 virus (Figure 4E). The genomic RNA/PFU ratio of R203/G204 virus was 1.6-2.3 folds higher than that of 203K/204R virus (Figure 4F), indicating that R203K/G204R increased the infectivity of SARS-CoV-2 produced from a human lung cell line.

**Figure 4.**
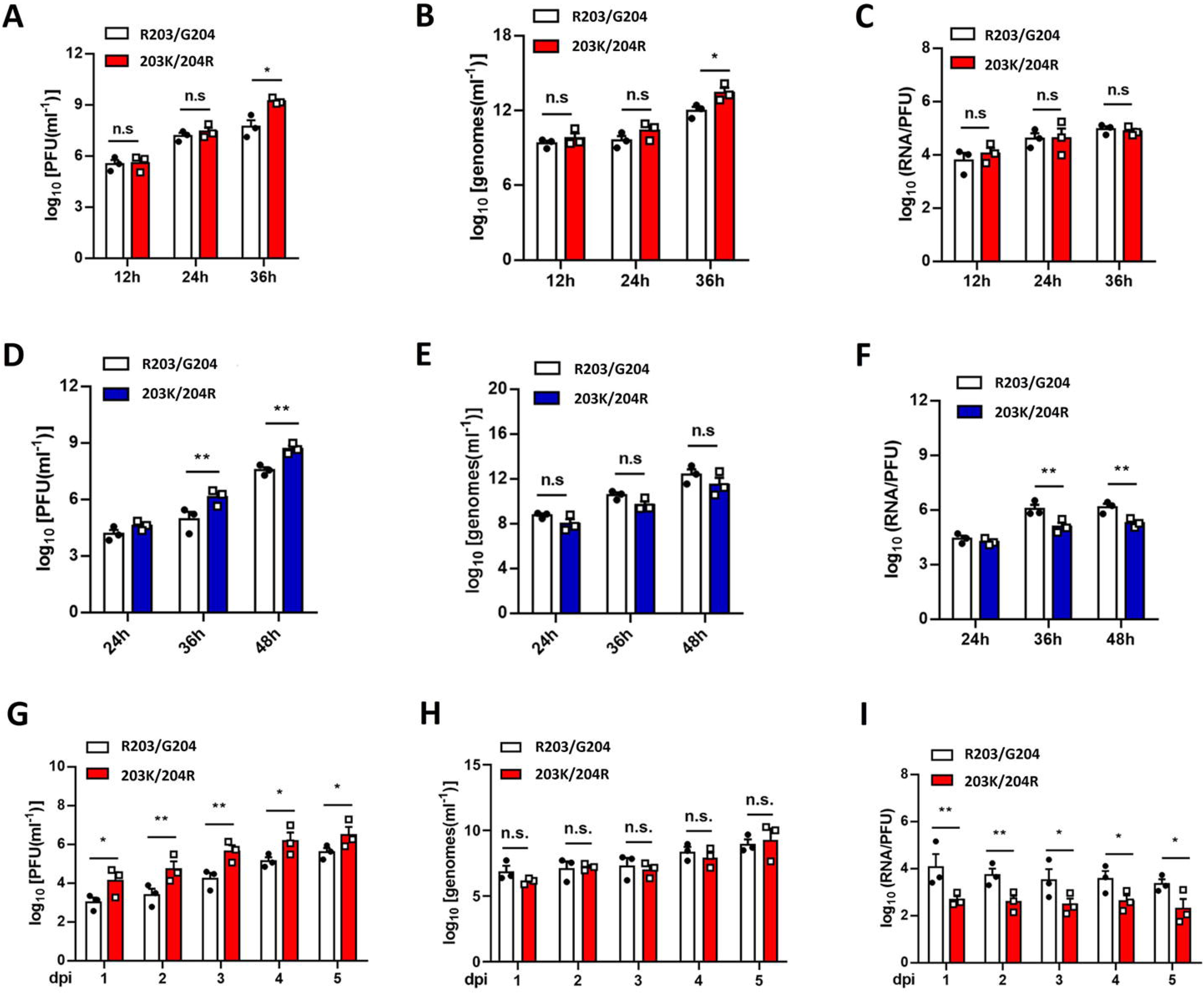
Effects of 203K/204R on viral replication and infectivity. (A-F) Viral replications and genomic RNA/PFU ratios of R203/G204 and 203K/204R viruses produced from Vero E6 (A-C) and Calu-3 (D-F) cell cultures. Cells were infected at a MOI of 0.01. Infectious viral titres (A, D) and genomic RNA levels (B, E) in the culture medium were determined by plaque assay and qRT-PCR, respectively. The genomic RNA/PFU ratios (C, F) were calculated to indicate virion infectivity. Data were represented as mean±s.e.m.. *, p<0.05, **, p<0.01. (G-I) Viral replications and genomic RNA/PFU ratios of R203/G204 and 203K/204R viruses. R203/G204 and 203K/204R viruses were inoculated onto primary human airway tissues at a MOI of 5. After incubation for 2 h, the culture was washed with PBS and maintained for 5 days. Infectious viral titres (G) and genomic RNA levels (H) in the culture medium were determined by plaque assay and qRT-PCR, respectively. The genomic RNA/PFU ratios (I) were calculated to indicate virion infectivity. (G-I) show R203K/G204R substitution enhances SARS-CoV-2 replication in primary human airway tissues.

Thereafter, we performed a comparison in hamsters. Three- to four-week-old hamsters were infected intranasally with 2×10^4^ PFU of R203/G204 or 203K/204R viruses. Infected hamsters from both groups exhibited similar weight loss (Figure 5A). On the 4th day post-infection (dpi), infectious viral titres from nasal washes and trachea were consistently higher in the hamsters infected with 203K/204R virus compared with those infected with R203/G204 virus (Figure 5B). We further compared the infectivity of R203/G204 and 203K/204R viruses produced in hamsters by determining their viral RNA levels and viral RNA/PFU ratios. The two viruses produced nearly identical levels of viral RNA across all organs (Figure 5C). The RNA/PFU ratio of 203K/204R virus was significantly lower than that of R203/G204 virus in nasal wash on 4 dpi (Figure 5D).

**Figure 5.**
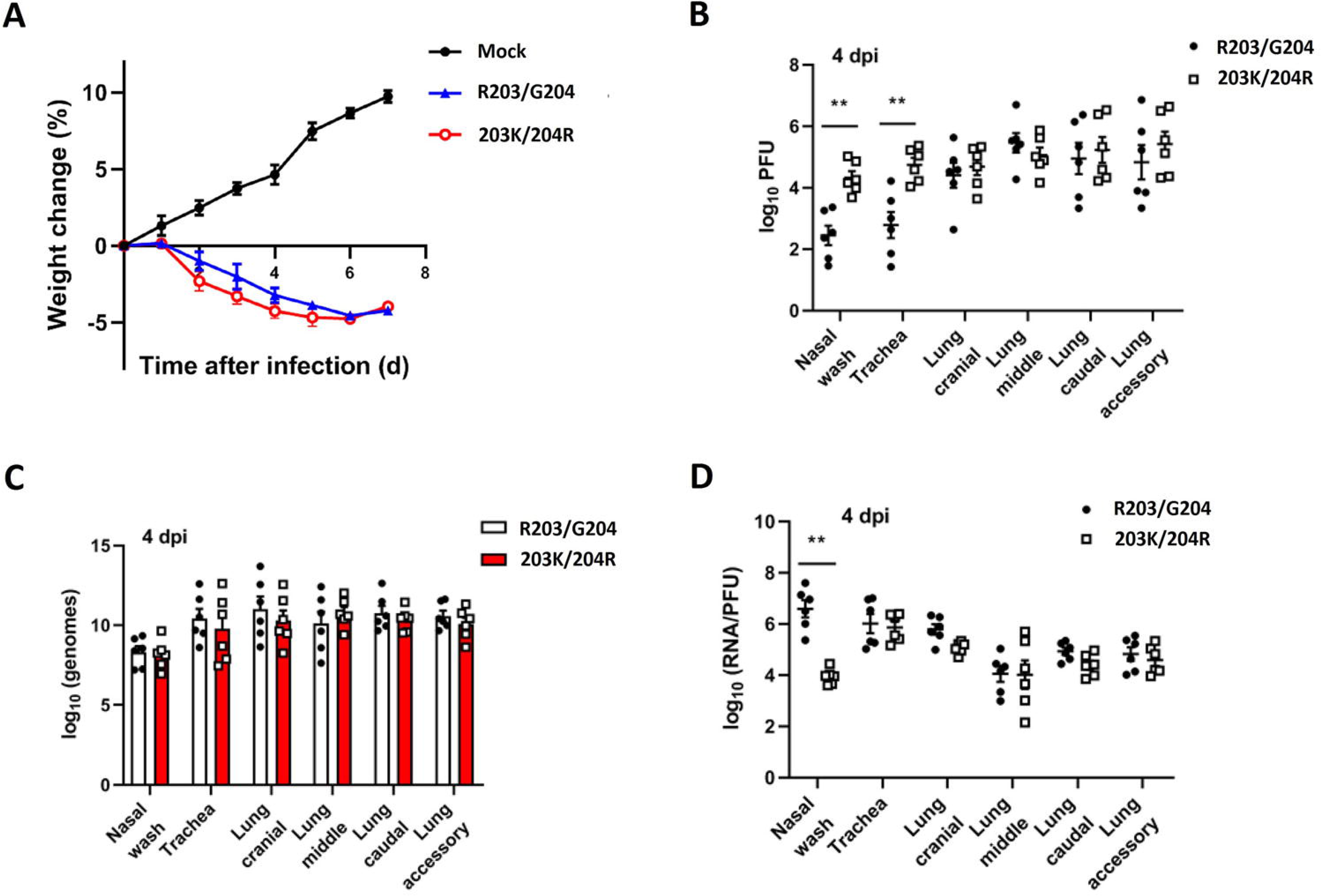
The infectivity of R203/G204 and 203K/204R viruses produced in hamsters. (A-D) Three- to four-week-old hamsters were infected intranasally with 2×10^4^ PFU of R203/G204 or 203K/204R viruses or PBS (mock). All data were from a single experiment. Weight loss (A) was monitored for 7 days post-infection. Infectious titres (B) and amounts of viral genomes (C) were measured in the nasal wash, trachea and lung on the 4th day post-infection (dpi). Genome/PFU ratios (D) on 4 dpi were calculated as a measure of infectivity. Data were represented as mean±s.e.m.. **, p<0.01.

In the comparison in a human airway model, we found that the infectious viral titres of 203K/204R virus were significantly higher than those of R203/G204 virus (Figure 4G). By contrast, there were no differences in viral RNA yields between the variants (Figure 4H). The genomic RNA/PFU ratios of R203/G204 virus were higher than those of 203K/204R virus (Figure 4I). These results demonstrated that the R203K/G204R mutation enhanced viral replication through increasing virion infectivity in primary human upper airway tissues. In consistent to our experimental results, we previously observed a negative correlation of R203/G204 IF with the cycle threshold for positive signal in E gene-based RT-PCR (Ct)^10^.

We measured the neutralization titres of a panel of sera collected from hamsters that was previously infected with R203/G204 virus. Each serum was analyzed using mNeonGreen reporter R203/G204 or 203K/204R viruses. All sera exhibited 1.14- to 2.08-fold higher neutralization titres (mean 1.51-fold) against heterologous 203K/204R virus compared with the homologous R203/G204 virus (Figure S14A-I). Serum 3 represented the highest neutralization titre (Figure S14C). The results suggested that R203K/G204R might confer a higher susceptibility to serum neutralization.

### 203K/204R virus show an association with the increased severity of disease

In experiments, we observed that the hamsters infected with 203K/204R virus had more extensive inflammatory damage with blood vessel congestion in lungs than R203/G204 (Figure 6A, B). For a further validation, we performed statistics analyses of sequenced strains with patient information (Table S5, S6, from GISAID). We found 203K/204R virus show a significant increase in the ratios of symptomatic : asymptomatic, hospitalized : outpatient, mild : severe and deceased : released (Figure 6C-F). For a prevention of the bias resulted from geological difference, we performed statistics of the data collected in a small scale (Figure 6C-F, S15A-D). Through correlation analyses between the IF of 203K/204R and the case fatality rate (CFR) per month in countries (Table S7), we found a significant positive correlation between the IF of 203K/204R and CFR (Figure 6G-I). These confirmed an increased virulence of 203K/204R.

**Figure 6.**
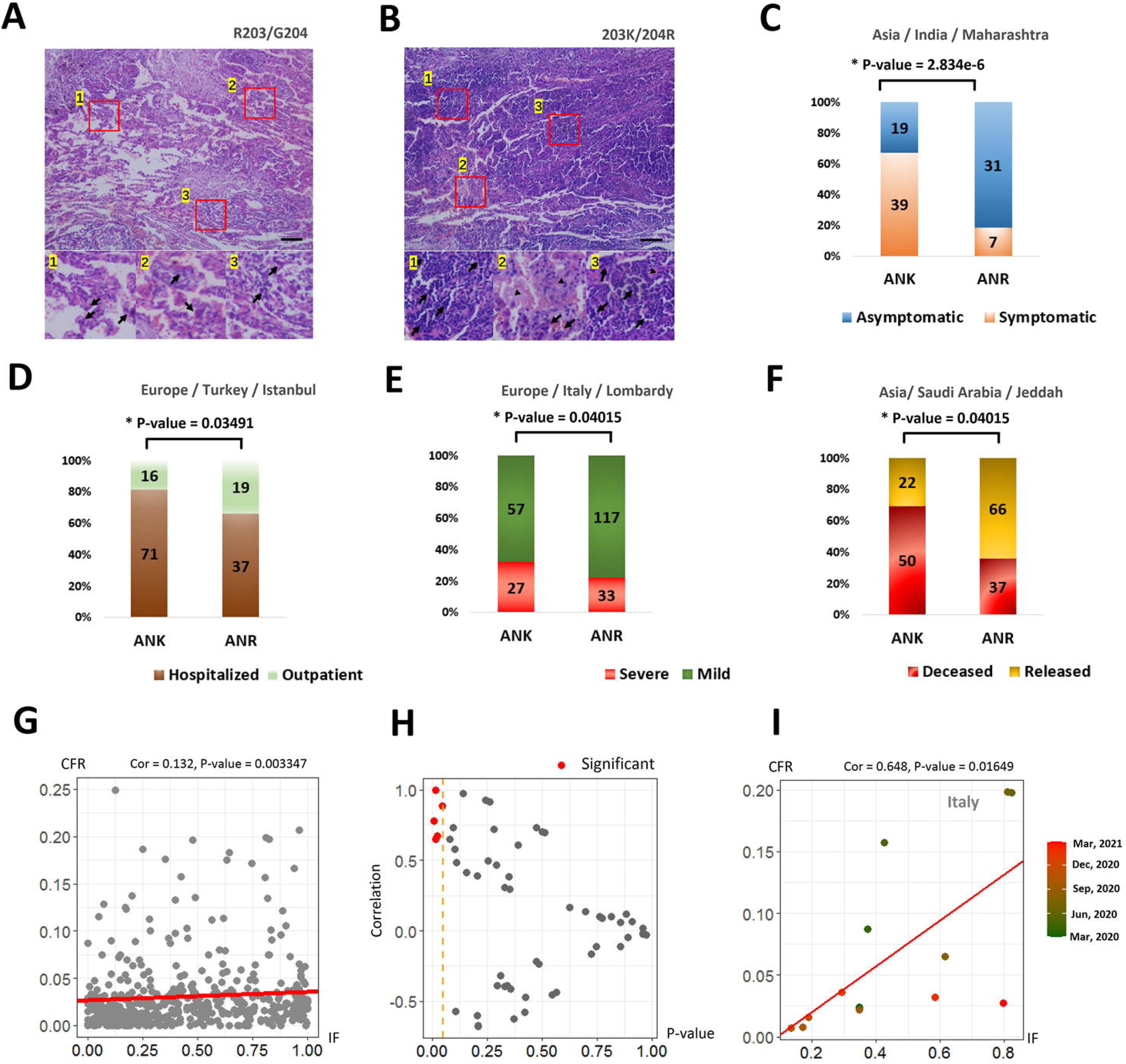
Effects of R203K/G204R on the severity of disease. (A,B) Severity between samples infected by R203/G204 and 203K/204R. Haemotoxylin and eosin (H&E) staining of lung sections from R203/G204 and 203K/204R viruses infected hamsters, collected at 7 days post-infection. The lower photographs depicted higher magnification images of the regions denoted by rectangles in upper photographs. Upper panel showed inflammatory damage with blood vessel congestion. Lower panel showed bronchiolar with aggregation of inflammatory cells (arrow) and surrounding alveolar wall infiltration (arrowhead). Scale bar = 100 μm. (C-F) Prediction on the clinical outcomes of ANR (R203/G204) and ANK (203K/204R). Four pairs of colors denote four pairs of opposite patient status. Collection places are listed at the top of figures. Y axis are the ratios between lineages with opposite patient status. Lineage numbers are marked. We tested the significance of ratio change by Chi-square Test. (G-I) Correlation analysis results between mutant IF and CFR. (G) Correlation analysis results between the ratios of mutants (x-axis) and the median case fatality rate (CFR) of COVID-19 in months and in countries (y-axis). The P-value was calculated by Spearman's test. (H) Scatter map showing the distribution of the IF-CFR correlation (Y-axis) and the P-value (X-axis) in countries. A vertical orange dotted line denotes the 0.05 P-value cutoff. (I) A specific example showing the correlation of mutant IFs and CFRs in one country, Italy. Continuous changing colors denote the months of IF-CFR pairs.

### Structural implications of R203K/G204R changes based on prediction

To investigate the impact of R203K/G204R onto the tertiary structure of the nucelocapsid, we built a tentative model of the N protein based on documented cryo-EM reports ^31^ (Figure 7). We found the N-terminal (NTD) and C-terminal domain (CTD) were docked in Murine hepatitis virus (MHV) nucleocapsid density^31^ in a reverse L-shape manner (Figure 7A). According to a previous study ^32^, the CTD dimer was suggested to be an assembly unit of N. The R203K/G204R mutation takes place in the linker region of N. The linker region is a serine/arginine-rich region with high flexibility. Due to the abundance of arginine, the linker region is more alkaline than NTD and CTD (Figuer 7B), i.e. the PI of NTD and CTD is around 10, while the PI of the linker region reaches 11.88. As lysine is also a basic amino acid, the R203K mutation only causes a small change in the PI of the linker region. But in contrast, the G204R introduce an extra basic amino acid leading to a more significant rise in linker region (PI). The electrostatic surface potential of N shows that the CTD dimer, also known as RNA-binding dimerization domain, is rich in positive charge (Figure 7C). However, owing to the lack of available 3D structure, the existing surface electrostatic potential analyses ignore the linker region. The linker region is highly basic, i.e. positive charged, which suggests that this region is also involved in the RNA binding process. Thus, R203K/G204R possibly impacts virus assembly.

**Figure 7.**
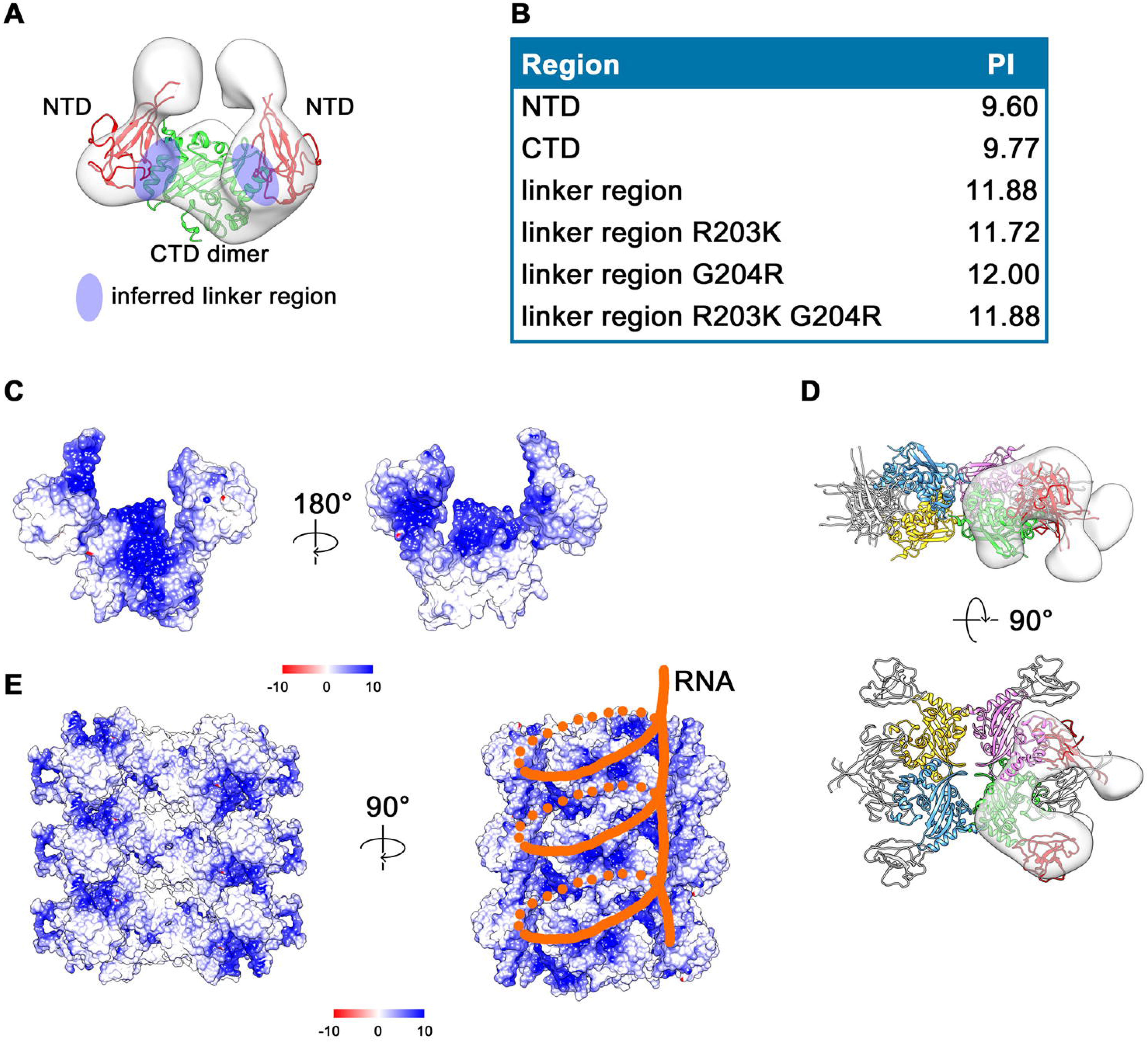
Tentative Model of the SARS-CoV-2 nucleocapsid. (A) Docking of the crystal structures of the SARS-CoV-2 nucleocapsid protein NTD monomer (PDB ID: 6M3M) and CTD dimer (PDB ID: 6WZQ) into the 3D density map of MHV nucleocapsid. The NTDs are colored red. The CTD dimer is colored green. The inferred linker region was highlighted in blue. The theoretical PI of each region is shown in (B). The electrostatic surface potential corresponding to (A) is shown in (C). (D) Four nucleocapsid protein dimers package in an asymmetry unit of the nucleocapsid. CTD dimers are colored blue, yellow, magneta and green, respectively. Density map of one asymmetry subunit is shown in gray semi-transparent surface with the corresponding fitted ribbon models colored green (CTDs) and red (NTDs). NTDs of other three nucleocapsid protein dimers are colored gray. Based on the model proposed in (C) and (D), we built a tentative model of nucleocapsid (E). In D and E, positively charged region in the surface is colored blue, as shown in the scale bar, −10 to 10 Kcal/(mol·e-). The possible RNA binding grooves is indicated by orange lines.

A more complete nucleocapsid model was proposed also based on cryo-EM reports ^31^. Four N dimers pack head to head forming an “X” shape (Figure 8D), and then packaging into a helical filament (Figure 7E). The CTD dimer form the core of the helical nucleocapsid, and the NTDs form the two arms outside the building block. Based on the surface electrostatic potential of the proposed model, the possible RNA binding groove of the helical nucleocapsid model is identified (Figure 7E). According to the winding path of the RNA, the RNA also surrounds and possibly interacts with linker region between the core (CTD) and the arms (NTD). Combining with the fact mentioned above that the R203K/G204R mutations could impact the local charge of the N protein, the R203K/G204R mutations may promote the binding of RNA by increasing the positive charge within the linker region, thereby accelerating the replication process.

## Discussions

In our previous work, we observed the rapid spread of R203K/G204R in the initial four months after the pandemic of SARS-CoV-2 and predicted that the mutants possibly benefit virus replication based on statistics ^10^. In this study, we not only validated the increase of infectivity by experiments, but also identified and validated the increase of virulence and fitness. We found the increase in infectivity is due to the promotion of virus replication, which may be caused by the change in the local charge of the N protein resulted from R203K/G204R. The increase in replication may lead to the increase of virulence and fitness in the end. Other high infectivity related mutations, such as D614G and N510Y, also showed association with the increased fitness and severity of diseases ^9,10,33–35^. R203K/G204R is akin to predominance like D614G (Figure 1) and is shared by rapidly growing lineages, B.1.1.7 and P.1. Understanding the effects of this mutation should be important for a further prevention of SARS-CoV-2.

Since the pandemic of SARS-CoV-2, more attentions are focused on S mutations. Out findings indicated that N mutations also altered the function and fitness of virus. Although S protein and N protein locate on the different part of a virion, there are a lot of similarities between D614G and R203K/G204R related to the consequences of the virus property. R203K/G204R variants also have an increased infectivity and fitness and an association with the severity of disease. The mutant 614G is reported to confer a higher neutralization titers than D614 ^9^. 203K/204R viruses, are also more susceptible to neutralization than R203/G204 viruses. In the identified 96 mutations, 15 (15.6%) are N mutations while 14 (14.6%) are S mutations (Table S2). For B.1.1.7, one quarter (7/28) are N mutations and one quarter (7/28) are S mutations (Table S4). The N mutation should be considered equally important in the future surveillance.

Although 203K/204R variants have a higher fitness than the preceding R203/G204 variants, the IF of 203K/204R did not increase continuously like 614G. The reason may be the cooperation and competition between 203K/204R and other mutants. We did not observed positive signatures for 203K/204R in some months and countries (Figure 3A, S13), these may be due to the effects of nearby adaptive mutants. Possibly for the same reason, we observed positive selection signatures for R203/G204 variants. The lineage ANK is oppressed by VNR, and then VNR is oppressed by AYK. We observed the existence of a certain amount of 501Y and 222V in the first half year of 2020 form the intra-host analysis results (Figure S3), indicating that the locations of N510Y (LG_5) and A222V (LG_4) are possibly hot points for mutating.

203K/204R, 501Y and 222V show higher fitness than R203/G204, N501 and A222, respectively. However, the identified frequencies of VNK, VYK and VYR are nearly zero in countries over time (Figure S8). The causation possibly is the antagonism of functional effects between A222V and another two mutations, R203K/G204R and N501Y. There is a negative correlation between 222V and the CFR but a positive correlation between 501Y and the CFR (Figure S15E-J, Table S8, S9). Previous efforts also inferred that N501Y has potentials to increase the fatality risk ^33^. R203K/G204R variants are identified with increased virulence as described above. The co-occurring of 222V and the other two mutants may counter-act each other in effects, thus the newly evolved lineages possibly are not more advantageous than the preceding variants. However, with the change of intra- and extra- environment over time, the antagonism may vanish at some time. R203K/G204R and N501Y both have increased infectivity and virulence. The combination of 203K/204R, 501Y and other mutants in LG_5 is B.1.1.7, which is reported to have an increased transmission ^25^ and be in association with the increased mortality of disease ^26^. 501Y alone with 203K/204R and other mutants co-occur in P.1, which also show an association with the severity of disease ^36,37^. The increase in infectivity, virulence of R203K/G204R variants should contributed to the effects referred above. R203K/G204R showed a high susceptibility to neutralization, thus the expansion of this mutation should not be obstacles for vaccines. R203K/G204R did not make contribution in the resistance to neutralization to B.1.1.7 and P.1. However, B.1.1.7 and P.1 have immune evasion capability^38,39^ for carrying neutralization-resistant mutants, such as 501Y and 484K. A recently evolved Indian lineage B.1.617 also carried a point mutation in 28881, but the mutation is G28881T not G28881A ^40^. Meanwhile, there is no mutation in 28882 and 28883. B.1.617 has a novel N mutation, R203M, instead of R203K/G204R. This novel mutation at the same location of R203K/G204R may enforce novel functional effects, inferring the importance of 28881-28883 in the SARS-CoV-2 genome.

## Methods

### Identification and statistics of the SARS-CoV-2 mutations

The full-length 884736 SARS-CoV-2 genomes are downloaded from GISAID (www.gisaid.org), NCBI and CoVdb ^41^. The collection dates of these samples ranged from December, 2019 to March 2021. We assumed the sequence of the strain MN908947, collected in December 2019^42^, as the ancestral state of the SARS-CoV-2 genome and performed genomic alignments between the 884736 SARS-CoV-2 strains and MN908947 by MUSCLE ^43,44^. Based on these alignments, we identified mutations and performed statistics of the monthly incidence frequency (IF) by a serious of Perl Scripts used before ^10^. We put samples collected in Dec-2019 and in Jan-2020 in a month Jan-2020 for convenience in statistics. We filtered out mutations with an IF in one month higher than 0.05 and finally identified 96 mutations. Next, we performed linkage disequilibrium analysis of these mutations, according to published algorithms ^45–47^. We calculated ρ^2^ and LOD, which are a squared correlation coefficient ^46^, and a statistical test to infer the linkage of two loci ^45^, respectively. We put mutations with ρ^2^ > 0.7 into one linkage group. The cutoff is based on the distribution of all ρ^2^ (Figure S2). We finally identified 12 mutation linkage groups and 26 singletons. In accordance to previous findings, we also identified LG_1 (D614G), LG_2 (L84S) and LG_3 (R203K/G204R).

We performed intra-host single nucleotide variation (iSNV) analyses following previous efforts ^10,48–50^ onto the raw sequence data including 14108 downloadable samples from United Kingdom (PRJEB37886). We processed reads by Bowtie2 ^51^, GATK Mark Duplicates ^52,53^ and SAMtools ^54^, requiring the mapping quality > 20 ^54^.

### Evolutionary analysis

We build a multiple sequence alignment file through retrieving sequences from pairwise sequence alignments using MN908947 as the reference. We performed sliding window analysis with a window size 200 bp and a step size 50 bp. We calculated Pi, Theta ^55^ and Tajima’s D ^56^ by VariScan 2.0 ^57,58^, and the composite likelihood ratio (CLR, step size = 50) ^28,29^ by SweepFineder2 ^59^. To test if one mutation site is under positive selection, we compared the median of the CLR in the 100 bp nearby region of the mutation (m) and the top 5% threshold (t) of all CLR values. We used CLR _m/c_ = m/c to infer the significance of peaks. A CLR _m/c_ > 1 imply a significant CLR peak and a positive selection evidence. We calculated the CLR _m/t_ in a 3000 bp region centering 28881-28883, the location of R203K/G204R. We performed the calculation for R203/G204 and 203K/204R variants repectively, and them made a comparison.

We extracted the nucleotides at the 96 mutation sites from all SARS-CoV-2 genomes and assembled the nucleotides into consecutive sequences. In other words, we used a shortened sequence (SS) with full mutaion information to represent the the whole genome. We clusterred these SS by month and by allele. For comparisons of Pi, Theta and Tajima’s D between R203/G204 and 203K/204R, we used the BioPerl PopGen library ^60^ to perform population genetic calculation within R203/G204 SS and 203K/204R SS, respectively.

We used the SS to construct the phylogenetic tree of SARS-CoV-2. For a better clarification of the relationship between strains in the phylogenetic tree, we discarded duplicate SS in the same collection month and collection country. We finally made a unique SS dataset with 322893 sequences. We built a multiple alignment file based on the alignments of these SS and the reference. Then we constructed the phylogenetic tree by FastTree ^61^. We used a R script library ggtree ^62^ to mark and color the phylogenetic tree. Marking and coloring are according to lineages, collection months or collection regions. We used the software GIF Movie Gear (www.gamani.com) to build animations for a show of the IF change in countries over time.

### Function prediction based on clinical and epidemic data

We manually gathered the clinical information of 36750 SARS-CoV-2 strains from GISAID. Like we did before ^10^, we grouped these information into four pairs of opposite patient status by a serious of keywords (Table S5). We counted the ratios of variants with different patient status and tested the significance by Chisq-test.

We calculated the case fatality rates (CFR) in countries using the epidemic data provided by ourworldindata.org/covid-deaths. We performed the correlation test between the median CFR and the IF of mutations in months. For the mean duration from the onset of symptoms to death of COVID-19 is 17 or 18 days ^63^, we performed an x+1 correlation test, in which an IF in month x is compared with a CFR in month x+1. We evaluated the significance of correlation through a two sides test.

### Generation of R203K/G204R mutant virus

The SARS-CoV-2 virus was generated by using a reverse genetic method ^9,64,65^. Seven different DNA fragments spanning the entire genome of SARS-CoV-2 (USA_WA1/2020 SARS-CoV-2 sequence, GenBank accession No. MT020880) were synthesized by Beijing Genomics Institute (BGI, Shanghai, China) and cloned into pUC57 or pCC1 (kindly provided by Dr. Yonghui Zheng) plasmids by standard molecular cloning methods. The sequences of F1~F7 fragments and the restriction enzymes used for digestion and ligation were listed in Supplementary information (Figure S16). The assembly of full-length cDNA and recovery of recombinant SARS-CoV-2 virus were performed as previously described ^9,64,65^. Briefly, the full-length cDNA of SARS-CoV-2 were assembled by in vitro ligation of contiguous cDNA fragments. Then full-length genomic RNA was collected by in vitro transcription and electroporated into Vero E6 cells. SARS-CoV-2 virus was harvested at 40 h post-electroporation and viral titers were determined by plaque assay. For the generation of R203K/G204R mutant virus, nucleotide substitutions of GGG→AAC were introduced into a subclone of pUC57-F7 containing the nucleocapsid gene of the SARS-CoV-2 wild-type infectious clone by overlap-extension PCR. The primers are shown in Figure S17.

### Animals, cells and infection

Three- to four-week-old hamsters were infected intranasally with 2×10^4^ PFU of R203/G204 or 203K/204R viruses. African green monkey kidney epithelial Vero E6 cells (ATCC) and Human lung adenocarcinoma epithelial Calu-3 cells (ATCC) were maintained in a high-glucose Dulbecco’s modified Eagle’s medium (DMEM, Gibco) supplemented with 10% FBS (Gibco) at 37⍰°C with 5% CO_2_. Cells were infected at a MOI of 0.01 for indicated time. All experiments with live virus were performed under biosafety level 3 (BSL3+) conditions.

### Plaque assay

Approximately 1×10^6^ cells were seeded to each well of 6-well plates and cultured at 37⍰°C, 5% CO_2_ for 12 h. R203/G204 or 203K/204R viruses were serially diluted in either DMEM with 2% FBS and 200 μL was transferred to the monolayers. The viruses were incubated with the cells for 1 h. After the incubation, overlay medium was added to the infected cells per well. The overlay medium contained DMEM with 2% FBS and 1% sea-plaque agarose. After 2 days incubation, plates were stained with neutral red and plaques were counted on a light box.

### Neutralization assay

Neutralization assays were performed using R203/G204 and 203K/204R mNeonGreen viruses as previously described ^9^. In brief, Vero cells were plated in 96-well plates. On the following day, sera were serially diluted and incubated with R203/G204 or 203K/204R mNeonGreen viruses at 37⍰°C for 1 h. The virus-serum mixture was transferred to the Vero cell plate with the final MOI of 2.0. After 20 h, Hoechst 33342 solution was added to stain cell nucleus, sealed with membrane, incubated at 37⍰°C for 20 min, and quantified for mNeonGreen fluorescence. The total cells (indicated by nucleus staining) and mNeonGreen-positive cells were quantified for each well. Infection rates were determined by dividing the mNeonGreen-positive cell number to total cell number. Relative infection rates were obtained by normalizing the infection rates of serum-treated groups to those of non-serum-treated controls. A nonlinear regression method was used to determine the dilution fold that neutralized 50% of mNeonGreen fluorescence (NT_50_). The curves of the relative infection rates versus the serum dilutions (log_10_ values) were plotted using GraphPad Prism 8.

### Viral infection in a primary human airway tissue model

For viral replication kinetics, either R203/G204 or 203K/204R virus was inoculated onto the culture at a MOI of 5 in PBS. After 2 h infection at 37⍰°C with 5% CO2, the inoculum was removed, and the culture was washed three times with PBS. The infected epithelial cells were maintained without any medium in the apical well, and medium was provided to the culture through the basal well. The infected cells were incubated at 37⍰°C, 5% CO2. From day 1 to day 5, 300 μL PBS was added onto the apical side of the airway culture and incubated at 37⍰°C for 30 min to elute the released viruses.

### Validation of competition assay

The R203/G204 and 203K/204R viruses were mixed at ratios of 1:1, 3:1 and 9:1. After competition assay, the total RNAs of these mixed viruses were isolated and amplified by RT-PCR. The ratios of 203K/204R to R203/G204 viruses were calculated by the peak heights of Sanger sequencing.

### 3D structure prediction of nucleocapsid protein and structural analyses

Atomic models (PDB accession code 6M3M, 6WZQ) of the nucleocapsid protein N-terminal domain (NTD) and C-terminal domain (CTD) were fitted to the Murine hepatitis virus (MHV) RNP densities^31^ using the ‘fit to segments’ tool in UCSF Chimera ^31^. The theoretical isoelectric point (pI) of each region was predicted by the Compute pI/Mw tool on the ExPASy Server ^31^. The electrostatic surface potential was generated with APBS ^31^ and viewed in UCSF Chimera.

### Haplotype network

We first created 7-bp sequences by joining the nucleotides of strains in seven mutation sites (R203K/G204R (LG_3), N501Y (LG_5), A222V (LG_4), C313T, S477N, S194L and I120F). We then calculated the IF and geological distribution of these 7-bp sequences. With this data, we plotted the haplotype network by PopArt ^66^.

### Others

We used the R package “corrplot” to perform correlation tests between pairs of LGs/mutations. We wrote Perl scripts to classify strains into lineages and counted the IFs of these lineages. Heatmaps, box-plots and scatter-plots were generated using the R libraries “gdata” and “ggplot2”.

## Supporting information

Supplementary Animations

## Author contributions

Z.Z., G.M, H.W., K.M., Y.X., G.L., P.D. and H.L. collected the data and performed population genetic analyses. N.X., H.W., B.F., W.X. and X.L. performed the experiments. G.M., Z.Z, K.M. and W.Z. performed the protein structure analysis. Z.Z. and G.M. conceived the idea. Z.Z., G.M. and X.L. wrote the manuscript. Z.Z., X.L. and G.M. coordinated the project.

## Acknowledgments

We gratefully acknowledge the submitting and the originating laboratories where genetic sequence data were generated and shared via NCBI and the GISAID Initiative. This work was supported by grants from the National Natural Science Foundation of China, SGC’s Rapid Response Funding for COVID-19 (C-0002), the National Key Research and Development Program (2019YFC1604600), the National Natural Science Foundation of China (81970008 and 31200941), the Fundamental Research Funds for the Central Universities (106112016CDJXY290002), the National Natural Science Foundation of HeBei province (19226631D).

## Notes

### Competing Interest Statement

The authors have declared no competing interest.

