## Supplementary Figure S16 for "Nucleocapsid mutation R203K/G204R increases the infectivity, fitness and virulence of SARS-CoV-2"

pUC57-F1 Bsal yellow shades indicate recognition sites

GGTCTCAtgatccgctaatacgactcactatagattaaaggtttataccttcccaggtaacaaaccaaccaactttcgatctcttgtagatc
tgttctctaaacgaactttaaaatctgtgtggctgtcactcggctgcatgcttagtgcactcacgcagtataattaataactaattactgtcgttg
acaggacacgagtaactcgtctatcttctgcaggctgcttacggtttcgtccgtgttgcagccgatcatcagcacatctaggtttcgtccgggtg
tgaccgaaaggtaagatggagagccttgtccctggtttcaacgagaaaacacacgtccaactcagtttgcctgttttacaggttcgcgacgtg
ctcgtacgtggctttggagactccgtggaggaggtcttatcagaggcacgtcaacatcttaaagatggcacttgtggcttagtagaagttgaa
aaaggcgttttgcctcaacttgaacagccctatgtgttcatcaaacgttcggatgctcgaactgcacctcatggtcatgttatggttgagctggt
agcagaactcgaaggcattcagtacggtcgtagtggtgagacacttggtgtccttgtccctcatgtgggcgaaataccagtggcttaccgcaa
ggttcttcttcgtaagaacggtaataaaggagctggtggccatagttacggcgccgatctaaagtcatttgacttaggcgacgagcttggcac
tgatccttatgaagattttcaagaaaactggaacactaaacatagcagtggtgttacccgtgaactcatgcgtgagcttaacggaggggcat
acactcgctatgtcgataacaacttctgtggccctgatggctaccctcttgagtgcattaaagaccttctagcacgtgctggtaaagcttcatgc
actttgtccgaacaactggactttattgacactaagaggggtgtatactgctgccgtgaacatgagcatgaaattgcttggtacacggaacgt
tctgaaaagagctatgaattgcagacaccttttgaaattaaattggcaaagaaatttgacaccttcaatggggaatgtccaaattttgtatttc
ccttaaattccataatcaagactattcaaccaagggttgaaaagaaaaagcttgatggctttatgggtagaattcgatctgtctatccagttgc
gtcaccaaatgaatgcaaccaaatgtgcctttcaactctcatgaagtgtgatcattgtggtgaaacttcatggcagacgggcgattttgttaaa
gccacttgcgaattttgtggcactgagaatttgactaaagaaggtgccactacttgtggttacttaccccaaaatgctgttgttaaaatttattg
tccagcatgtcacaattcagaagtaggacctgagcatagtcttgccgaataccataatgaatctggcttgaaaaccattcttcgtaagggtgg
tcgcactattgcctttggaggctgtgtgttctcttatgttggttgccataacaagtgtgcctattgggttccacgtgctagcgctaacataggttgt
aaccatacaggtgttgttggagaaggttccgaaggtcttaatgacaaccttcttgaaatactccaaaaagagaaagtcaacatcaatattgtt
ggtgactttaaacttaatgaagagatcgccattattttggcatctttttctgcttccacaagtgcttttgtggaaactgtgaaaggtttggattata
aagcattcaaacaaattgttgaatcctgtggtaattttaaagttacaaaaggaaaagctaaaaaaggtgcctggaatattggtgaacagaa
atcaatactgagtcctctttatgcatttgcatcagaggctgctcgtgttgtacgatcaattttctcccgcactcttgaaactgctcaaaattctgtg
cgtgttttacagaaggccgctataacaatactagatggaatttcacagtattcactgagactcattgatgctatgatgttcacatctgatttggct
actaacaatctagttgtaatggcctacattacaggtggtgttgttcagttgacttcgcagtggctaactaacatctttggcactgtttatgaaaa
actcaaacccgtccttgattggcttgaagagaagtttaaggaaggtgtagagtttcttagagacggttgggaaattgttaaatttatctcaacc
tgtgcttgtgaaattgtcggtggacaaattgtcacctgtgcaaaggaaattaaggagagtgttcagacattctttaagcttgtaaataaattttt
ggctttgtgtgctgactctatcattattggtggagctaaacttaaagccttgaatttaggtgaaacatttgtcacgcactcaaagggattgtaca
gaaagtgtgttaaatccagagaagaaactggcctactcatgcctctaaaagccccaaaagaaattatcttcttagagggagaaacacttccc
acagaagtgttaacagaggaagttgtcttgaaaactggtgatttacaaccattagaacaacctactagtgaagctgttgaagctccattggtt
ggtacaccagtttgtattaacgggcttatgttgctcgaaatcaaagacacagaaaagtactgtgcccttgcacctaatatgatggtaacaaac
aataccttcacactcaaaggcggtgcaccaacaaaggttacttttggtgatgacactgtgatagaagtgcaaggttacaagagtgtgaatat
cacttttgaacttgatgaaaggattgataaagtacttaatgagaagtgctctgcctatacagttgaactcggtacagaagtaaatgagttcgcc
tgtgttgtggcagatgctgtcataaaaactttgcaaccagtatctgaattacttacaccactgggcattgatttagatgagtggagtatggctac
atactacttatttgatgagtctggtgagtttaaattggcttcacatatgtattgttctttctaccctccagatgaggatgaagaagaaggtgattg
tgaagaagaagagtttgagccatcaactcaatatgagtatggtactgaagatgattaccaaggtaaacctttggaatttggtgccacttctgc
tgctcttcaacctgaagaagagcaagaagaagattggttagatgatgatagtcaacaaactgttggtcaacaagacggcagtgaggacaat
cagacaactactattcaaacaattgttgaggttcaacctcaattagagatggaacttacaccagttgttcagactattgaagtgaatagtttta
gtggttatttaaaacttactgacaatgtatacattaaaaatgcagacattgtggaagaagctaaaaaggtaaaaccaacagtggttgttaat
gcagccaatgtttaccttaaacatggaggaggtgttgcaggagccttaaataaggctactaacaatgccatgcaagttgaatctgatgattac
atagctactaatggaccacttaaagtgggtggtagttgtgttttaagcggacacaatcttgctaaacactgtcttcatgttgtcggcTGAGA
CC

pCC1-F2 BsaI

GGTCTCAcggcccaaatgttaacaaaggtgaagacattcaacttcttaagagtgcttatgaaaattttaatcagcacgaagttctac
ttgcaccattattatcagctggtatttttggtgctgaccctatacattctttaagagtttgtgtagatactgttcgcacaaatgtctacttag
ctgtctttgataaaaatctctatgacaaacttgtttcaagctttttggaaatgaagagtgaaaagcaagttgaacaaaagatcgctgag
attcctaaagaggaagttaagccatttataactgaaagtaaaccttcagttgaacagagaaaacaagatgataagaaaatcaaagct
tgtgttgaagaagttacaacaactctggaagaaactaagttcctcacagaaaacttgttactttatattgacattaatggcaatcttcat
ccagattctgccactcttgttagtgacattgacatcactttcttaaagaaagatgctccatatatagtgggtgatgttgttcaagagggtg
ttttaactgctgtggttatacctactaaaaaggctggtggcactactgaaatgctagcgaaagctttgagaaaagtgccaacagacaa
ttatataaccacttacccgggtcagggtttaaatggttacactgtagaggaggcaaagacagtgcttaaaaagtgtaaaagtgcctttt
acattctaccatctattatctctaatgagaagcaagaaattcttggaactgtttcttggaatttgcgagaaatgcttgcacatgcagaag
aaacacgcaaattaatgcctgtctgtgtggaaactaaagccatagtttcaactatacagcgtaaatataagggtattaaaatacaaga
gggtgtggttgattatggtgctagattttacttttacaccagtaaaacaactgtagcgtcacttatcaacacacttaacgatctaaatga
aactcttgttacaatgccacttggctatgtaacacatggcttaaatttggaagaagctgctcggtatatgagatctctcaaagtgccagc
tacagtttctgtttcttcacctgatgctgttacagcgtataatggttatcttacttcttcttctaaaacacctgaagaacattttattgaaac
catctcacttgctggttcctataaagattggtcctattctggacaatctacacaactaggtatagaatttcttaagagaggtgataaaag
tgtatattacactagtaatcctaccacattccacctagatggtgaagttatcacctttgacaatcttaagacacttctttctttgagagaag
tgaggactattaaggtgtttacaacagtagacaacattaacctccacacgcaagttgtggacatgtcaatgacatatggacaacagttt
ggtccaacttatttggatggagctgatgttactaaaataaaacctcataattcacatgaaggtaaaacattttatgttttacctaatgatg
acactctacgtgttgaggcttttgagtactaccacacaactgatcctagttttctgggtaggtacatgtcagcattaaatcacactaaaa
agtggaaatacccacaagttaatggtttaacttctattaaatgggcagataacaactgttatcttgccactgcattgttaacactccaac
aaatagagttgaagtttaatccacctgctctacaagatgcttattacagagcaagggctggtgaagctgctaacttttgtgcacttatct
tagcctactgtaataagacagtaggtgagttaggtgatgttagagaaacaatgagttacttgtttcaacatgccaatttagattcttgca
aaagagtcttgaacgtggtgtgtaaaacttgtggacaacagcagacaacccttaagggtgtagaagctgttatgtacatgggcacact
ttcttatgaacaatttaagaaaggtgttcagataccttgtacgtgtggtaaacaagctacaaaatatctagtacaacaggagtcacttt
tgttatgatgtcagcaccacctgctcagtatgaacttaagcatggtacatttacttgtgctagtgagtacactggtaattaccagtgtggt
cactataaacatataacttctaaagaaactttgtattgcatagacggtgctttacttacaaagtcctcagaatacaaaggtcctattacg
gatgttttctacaaagaaaacagttacacaacaaccataaaaccagttacttataaattggatggtgttgtttgtacagaaattgaccct
aagttggacaattattataagaaagacaattcttatttcacagagcaaccaattgatcttgtaccaaaccaaccatatccaaacgcaag
cttcgataattttaagtttgtatgtgataatatcaaatttgctgatgatttaaaccagttaactggttataagaaacctgcttcaagagag
cttaaagttacatttttccctgacttaaatggtgatgtggtggctattgattataaacactacacaccctcttttaagaaaggagctaaat
tgttacataaacctattgtttggcatgttaacaatgcaactaataaagccacgtataaaccaaatacctggtgtatacgttgtctttgga
gcacaaaaccagttgaaacatcaaattcgtttgatgtactgaagtcagaggacgcgcagggaatggataatcttgcctgcgaagatct
aaaaccagtctctgaagaagtagtggaaaatcctaccatacagaaagacgttcttgagtgtaatgtgaaaactaccgaagttgtagg
agacattatacttaaaccagcaaataatagtttaaaaattacagaagaggttggccacacagatctaatggctgcttatgtagacaatt
ctagtcttactattaagaaacctaatgaattatctagagtattaggtttgaaaacccttgctactcatggtttagctgctgttaatagtgtc
ccttgggatactatagctaattatgctaagccttttcttaacaaagttgttagtacaactactaacatagttacacggtgtttaaaccgtg
tttgtactaattatatgccttatttctttactttattgctacaattgtgtacttttactagaagtacaaattctagaattaaagcatctatgcc
gactactatagcaaagaatactgttaagagtgtcggtaaattttgtctagaggcttcatttaattatttgaagtcacctaatttttctaaac
tgataaatattataatttggtttttactattaagtgtttgcctaggttctttaatctactcaaccgctgctttaggtgttttaatgtctaattta
ggcatgccttcttactgtactggttacagagaaggctatttgaactctactaatgtcactattgcaacctactgtactggttctataccttg
tagtgtttgtcttagtggtttagattctttagacacctatccttctttagaaactatacaaattaccatttcatcttttaaatgggatttaact
gcttttggcttagttgcagagtggtttttggcatatattcttttcactaggtttttctatgtacttggattggctgcaatcatgcaattgttttt
cagctattttgcagtacattttattagtaattcttggcttatgtggttaataattaatcttgtacaaatggccccgatttcagctatggttag
aatgtacatcttctttgcatcattttattatgtatggaaaagttatgtgcatgttgtagacggttgtaattcatcaacttgtatgatgtgtta

caaacTGAGACC

pCC1-F3 Bsal

GGTCTCAaaacgtaatagagcaacaagagtcgaatgtacaactattgttaatggtgttagaaggtccttttatgtctatgctaatggaggt
aaaggcttttgcaaactacacaattggaattgtgttaattgtgatacattctgtgctggtagtacatttattagtgatgaagttgcgagagactt
gtcactacagtttaaaagaccaataaatcctactgaccagtcttcttacatcgttgatagtgttacagtgaagaatggttccatccatctttactt
tgataaagctggtcaaaagacttatgaaagacattctctctctcattttgttaacttagacaacctgagagctaataacactaaaggttcattg
cctattaatgttatagtttttgatggtaaatcaaaatgtgaagaatcatctgcaaaatcagcgtctgtttactacagtcagcttatgtgtcaacct
atactgttactagatcaggcattagtgtctgatgttggtgatagtgcggaagttgcagttaaaatgtttgatgcttacgttaatacgttttcatca
acttttaacgtaccaatggaaaaactcaaaacactagttgcaactgcagaagctgaacttgcaaagaatgtgtccttagacaatgtcttatct
acttttatttcagcagctcggcaagggtttgttgattcagatgtagaaactaaagatgttgttgaatgtcttaaattgtcacatcaatctgacata
gaagttactggcgatagttgtaataactatatgctcacctataacaaagttgaaaacatgacaccccgtgaccttggtgcttgtattgactgta
gtgcgcgtcatattaatgcgcaggtagcaaaaagtcacaacattgctttgatatggaacgttaaagatttcatgtcattgtctgaacaactacg
aaaacaaatacgtagtgctgctaaaaagaataacttaccttttaagttgacatgtgcaactactagacaagttgttaatgttgtaacaacaaa
gatagcacttaagggtggtaaaattgttaataattggttgaagcagttaattaaagttacacttgtgttcctttttgttgctgctattttctatttaa
taacacctgttcatgtcatgtctaaacatactgacttttcaagtgaaatcataggatacaaggctattgatggtggtgtcactcgtgacatagc
atctacagatacttgttttgctaacaaacatgctgattttgacacatggtttagccagcgtggtggtagttatactaatgacaaagcttgcccat
tgattgctgcagtcataacaagagaagtgggttttgtcgtgcctggtttgcctggcacgatattacgcacaactaatggtgactttttgcatttct
tacctagagtttttagtgcagttggtaacatctgttacacaccatcaaaacttatagagtacactgactttgcaacatcagcttgtgttttggctg
ctgaatgtacaatttttaaagatgcttctggtaagccagtaccatattgttatgataccaatgtactagaaggttctgttgcttatgaaagtttac
gccctgacacacgttatgtgctcatggatggctctattattcaatttcctaacacctaccttgaaggttctgttagagtggtaacaacttttgatt
ctgagtactgtaggcacggcacttgtgaaagatcagaagctggtgtttgtgtatctactagtggtagatgggtacttaacaatgattattacag
atctttaccaggagttttctgtggtgtagatgctgtaaatttacttactaatatgtttacaccactaattcaacctattggtgctttggacatatca
gcatctatagtagctggtggtattgtagctatcgtagtaacatgccttgcctactattttatgaggtttagaagagcttttggtgaatacagtcat
gtagttgcctttaatactttactattccttatgtcattcactgtactctgtttaacaccagtttactcattcttacctggtgtttattctgttatttactt
gtacttgacattttatcttactaatgatgtttctttttttagcacatattcagtggatggttatgttcacacctttagtacctttctggataacaattgc
ttatatcatttgtatttccacaaagcatttctattggttctttagtaattacctaaagagacgtgtagtctttaatggtgtttcctttagtacttttga
agaagctgcgctgtgcacctttttgttaaataaagaaatgtatctaaagttgcgtagtgatgtgctattacctcttacgcaatataatagatact
tagctctttataataagtacaagtattttagtggagcaatggatacaactagctacagagaagctgcttgttgtcatctcgcaaaggctctcaa
tgacttcagtaactcaggttctgatgttctttaccaaccaccacaaacctctatcacctcagctgttttgcagagtggttttagaaaaatggcatt
cccatctggtaaagttgagggttgtatggtacaagtaacttgtggtacaactacacttaacggtctttggcttgatgacgtagtttactgtccaa
gacatgtgatctgcacctctgaagacatgcttaaccctaattatgaagatttactcattcgtaagtctaatcataatttcttggtacaggctggta
atgttcaactcagggttattggacattctatgcaaaattgtgtacttaagcttaaggttgatacagccaatcctaagacacctaagtataagttt
gttcgcattcaaccaggacagactttttcagtgttagcttgttacaatggttcaccatctggtgtttaccaatgtgctatgaggcccaatttcact
attaagggttcattccttaatggttcatgtggtagtgttggttttaacatagattatgactgtgtctctttttgttacatgcaccatatggaattacc
aactggagttcatgctggcacagacttagaaggtaacttttatggaccttttgttgacaggcaaacagcacaagcagctggtacggacacaa
ctattacagttaatgttttagcttggttgtacgctgctgttataaatggagacaggtggtttctcaatcgatttaccacaactcttaatgactttaa
ccttgtggctatgaagtacaattatgaacctctaacacaagaccatgttgacatactaggacctctttctgctcaaactggaattgccgttttag
atatgtgtgcttcattaaaagaattactgcaaaatggtatgaatggacgtaccatattgggtagtgctttattagaagatgaatttacaccttttt
gatgttgttagacaatgctcaggtgttactttccaaagtgcagtgaaaagaacaatcaagggtacacaccactggttgttactcacaattttga
cttcacttttagttttagtccagagtactcaatggtctttgttctttttttgtatgaaaatgcctttttaccttttgctatgggtattattgctatgtct
gcttttgcaatgatgtttgtcaaacataagcatgcatttctctgtttgtttttgttaccttctcttgccactgtagcttattttaatatggtctatatgc

ctgctagttgggtgatgcgtattatgacatggttggatatggttgatactagtttgtctggttttaagctaaaagactgtgttatgtatgcatcag
ctgtagtgttactaatccttatgacagcaagaactgtgtatgatgatggtgctaggagagtgtggacacttatgaatgtcttgacactcgtttat
aaagtttattatggtaatgctttagatcaagccatttccatgtgggctcttataatctctgttacttctaactactcaggtgtagttacaactgtca
tgtttttggccagaggtattgtttttatgtgtgttgagtattgccctattttcttcataactggtaatacacttcagtgtataatgctagtttattgttt
cttaggctatttttgtacttgttactttggcctcttttgtttactcaaccgctactttagactgactcttggtgtttatgattacttagtttctacacag
gagtttagatatatgaattcacagggactactcccacccaagaatagcatagatgccttcaaactcaacattaaattgttgggtgttggtggca
aaccttgtatcaaagtagccactgtacagtctaaaatgtcagatgtaaagtgcacatcagtagtcttactctcagttttgcaacaactcagagt
agaatcatcatctaaattgtgggctcaatgtgtccagttacacaatgacattctcttagctaaagatactactgaagccttTGAGACC

pUC57-F4 BsaI

GGTCTCAcctttgaaaaaatggtttcactactttctgttttgctttccatgcagggtgctgtagacataaacaagctttgtgaagaaatgctg
gacaacagggcaaccttacaagctatagcctcagagtttagttcccttccatcatatgcagcttttgctactgctcaagaagcttatgagcagg
ctgttgctaatggtgattctgaagttgttcttaaaaagttgaagaagtctttgaatgtggctaaatctgaatttgaccgtgatgcagccatgcaa
cgtaagttggaaaagatggctgatcaagctatgacccaaatgtataaacaggctagatctgaggacaagagggcaaaagttactagtgcta
tgcagacaatgcttttcactatgcttagaaagttggataatgatgcactcaacaacattatcaacaatgcaagagatggttgtgttcccttgaa
cataatacctcttacaacagcagccaaactaatggttgtcataccagactataacacatataaaaatacgtgtgatggtacaacatttacttat
gcatcagcattgtgggaaatccaacaggttgtagatgcagatagtaaaattgttcaacttagtgaaattagtatggacaattcacctaatttag
catggcctcttattgtaacagctttaagggccaattctgctgtcaaattacagaataatgagcttagtcctgttgcactacgacagatgtcttgt
gctgccggtactacacaaactgcttgcactgatgacaatgcgttagcttactacaacacaacaaagggaggtaggtttgtacttgcactgtta
tccgatttacaggatttgaaatgggctagattccctaagagtgatggaactggtactatctatacagaactggaaccaccttgtaggtttgtta
cagacacacctaaaggtcctaaagtgaagtatttatactttattaaaggattaaacaacctaaatagaggtatggtacttggtagtttagctgc
cacagtacgtctacaagctggtaatgcaacagaagtgcctgccaattcaactgtattatctttctgtgcttttgctgtagatgctgctaaagctt
acaaagattatctagctagtgggggacaaccaatcactaattgtgttaagatgttgtgtacacacactggtactggtcaggcaataacagtta
caccggaagccaatatggatcaagaatcctttggtggtgcatcgtgttgtctgtactgccgttgccacatagatcatccaaatcctaaaggatt
ttgtgacttaaaaggtaagtatgtacaaatacctacaacttgtgctaatgaccctgtgggttttacacttaaaaacacagtctgtaccgtctgc
ggtatgtggaaaggttatggctgtagttgtgatcaactccgcgaacccatgcttcagtcagctgatgcacaatcgtttttaaacgggtttgcgg
tgtaagtgcagcccgtcttacaccgtgcggcacaggcactagtactgatgtcgtatacagggcttttgacatctacaatgataaagtagctggt
tttgctaaattcctaaaaactaattgttgtcgcttccaagaaaaggacgaagatgacaatttaattgattcttactttgtagttaagagacacac
tttctctaactaccaacatgaagaaacaatttataatttacttaaggattgtccagctgttgctaaacatgacttctttaagtttagaatagacg
gtgacatggtaccacatatatcacgtcaacgtcttactaaatacacaatggcagacctcgtctatgctttaaggcattttgatgaaggtaattgt
gacacattaaaagaaatacttgtcacatacaattgttgtgatgatgattatttcaataaaaaggactggtatgattttgtagaaaacccagata
tattacgcgtatacgccaacttaggtgaacgtgtacgccaagctttgttaaaaacagtacaattctgtgatgccatgcgaaatgctggtattgt
tggtgtactgacattagataatcaagatctcaatggtaactggtatgatttcggtgatttcatacaaaccacgccaggtagtggagttcctgttg
tagattcttattattcattgttaatgcctatattaaccttgaccagggctttaactgcagagtcacatgttgacactgacttaacaaagccttaca
ttaagtgggatttgttaaaatatgacttcacggaagagaggttaaaactctttgaccgttattttaaatattgggatcagacataccacccaaa
ttgtgttaactgtttggatgacagatgcattctgcattgtgcaaactttaatgttttattctctacagtgttcccacctacaagttttggaccacta
gtgagaaaaatatttgttgatggtgttccatttgtagtttcaactggataccacttcagagagctaggtgttgtacataatcaggatgtaaactt
acatagctctagacttagttttaaggaattacttgtgtatgctgctgaccctgctatgcacgctgcttctggtaatctattactagataaacgcac
tacgtgcttttcagtagctgcacttactaacaatgttgcttttcaaactgtcaaacccggtaattttaacaaagacttctatgactttgctgtgtct
aagggtttctttaaggaaggaagttctgttgaattaaaacacttcttctttgctcaggatggtaatgctgctatcagcgattatgactactatcgt
tataatctaccaacaatgtgtgatatcagacaactactatttgtagttgaagttgttgataagtactttgattgttacgatggtggctgtattaat
gctaaccaagtcatcgtcaacaacctagacaaatcagctggttttccatttaataaatggggtaaggctagactttattatgattcaatgagtt

atgaggatcaagatgcacttttcgcatatacaaaacgtaatgtcatccctactataactcaaatgaatcttaagtatgccattagtgcaaaga
atagagctcgcaccgtagctggtgtctctatctgtagtactatgaccaatagacagtttcatcaaaaattattgaaatcaatagccgccactag
aggagctactgtagtaattggaacaagcaaattctatggtggttggcacaacatgttaaaaactgtttatagtgatgtagaaaaccctcacct
tatgggttgggattatcctaaatgtgatagagccatgcctaacatgcttagaattatggcctcacttgttcttgctcgcaaacatacaacgtgtt
gtagcttgtcacaccgtttctatagattagctaatgagtgtgctcaagtattgagtgaaatggtcatgtgtggcggttcactatatgttaaacca
ggtggaacctcatcaggagatgccacaactgcttatgctaatagtgtttttaacatttgtcaagctgtcacggccaatgttaatgcacttttatct
actgatggtaacaaaattgccgataagtatgtccgcaatttacaacacagactttatgagtgtctctatagaaatagagatgttgacacagac
tttgtgaatgagttttacgcatatttgcgtaaacatttctcaatgatgatactctctgacgatgctgttgtgtgtttcaatagcacttatgcatctca
aggtctagtggctagcataaagaactttaagtcagttctttattatcaaaacaatgtttttatgtctgaagcaaaatgttggactgagactgacc
ttactaaaggacctcatgaattttgctctcaacatacaatgctagttaaacagggtgatgattatgtgtaccttccttacccagatccatcaaga
atcctaggggccggctgttttgtagatgatatcgtaaaaacagatggtacacttatgattgaacggttcgtgtctttagctatagatgcttaccc
acttactaaacatcctaatcaggagtatgctgatgtctttcatttgtacttacaatacataagaaagctacatgatgagttaacaggacacatg
ttagacatgtattctgttatgcttactaatgataacacttcaaggtattgggaacctgagttttatgaggctatgtacacaccgcatacagtctta
caggctgttggggcttgtgttctttgcaattcacagacttcattaagatgtggtgcttgcatacgtagaccattcttatgttgtaaatgctgttacg
accatgtcatatcaacatcacataaattagtcttgtctgttaatccgtatgtttgcaatgctccaggttgtgatgtcacagatgtgactcaacttt
acttaggaggtatgagctattattgtaaatcacataaaccacccattagttttccattgtgtgctaatggacaagtttttggtttatataaaaata
catgtgttggtagcgataatgttactgactttaatgcaattgcaacatgtgactggacaaatgctggtgattacattttagctaacacctgtact
gaaagactcaagctttttgcagcagaaacgctcaaagctactgaggagacatttaaactgtcttatggtattgctactgtacgtgaagtgctgt
ctgacagagaattacatctttcatgggaagttggtaaacctagaccaccacttaaccgaaattatgtctttactggttatcgtgtaactaaaaa
cagtaaagtacaaataggagagtacacctttgaaaaaggtgactatggtgatgctgttgtttaccgaggtacaacaacttacaaattaaatgt
tggtgattattttgtgctgacatcacatacagtaatgccattaagtgcacctacactagtgccacaagagcactatgttagaattactggcttat
acccaacactcaatatctcagatgagttttctagcaatgttgcaaattatcaaaaggttggtatgcaaaagtattctacactccagggaccacc
tggtactggtaagagtcattttgctattggcctagctctctactacccttctgctcgcatagtgtatacagcttgctctcatgccgctgttgatgca
ctatgtgagaaggcattaaaatatttgcctatagataaatgtagtagaattatacctgcacgtgctcgtgtagagtgttttgataaattcaaagt
gaattcaacattagaacagtatgtcttttgtactgtaaatgcattgcctgagacgacagcagatatagttgtctttgatgaaatttcaatggcca
caaattatgatttgagtgttgtcaatgccagattacgtgctaagcactatgtgtacattggcgaccctgctcaattacctgcaccacgcacattg
ctaactaagggcacactagaaccagaatatttcaattcagtgtgtagacttatgaaaactataggtccagacatgttcctcggaacttgtcgg
cgttgtcctgctgaaattgttgacactgtgagtTGAGACC

pUC57-F5 Esp3I
CGTCTCAgagtgctttggtttatgataataagcttaaagcacataaagacaaatcagctcaatgctttaaaatgttttataagggtgttatc
acgcatgatgtttcatctgcaattaacaggccacaaataggcgtggtaagagaattccttacacgtaaccctgcttggagaaaagctgtcttt
atttcaccttataattcacagaatgctgtagcctcaaagattttgggactaccaactcaaactgttgattcatcacagggctcagaatatgact
atgtcatattcactcaaaccactgaaacagctcactcttgtaatgtaaacagatttaatgttgctattaccagagcaaaagtaggcatactttg
cataatgtctgatagagacctttatgacaagttgcaatttacaagtcttgaaattccacgtaggaatgtggcaactttacaagctgaaaatgta
acaggactctttaaagattgtagtaaggtaatcactgggttacatcctacacaggcacctacacacctcagtgttgacactaaattcaaaact
gaaggtttatgtgttgacatacctggcatacctaaggacatgacctatagaagactcatctctatgatgggttttaaaatgaattatcaagtta
atggttaccctaacatgtttatcacccgcgaagaagctataagacatgtacgtgcatggattggcttcgatgtcgaggggtgtcatgctactag
agaagctgttggtaccaatttacctttacagctaggtttttctacaggtgttaacctagttgctgtacctacaggttatgttgatacacctaataa
tacagatttttccagagttagtgctaaaccaccgcctggagatcaatttaaacacctcataccacttatgtacaaaggacttccttggaatgta
gtgcgtataaagattgtacaaatgttaagtgacacacttaaaaatctctctgacagagtcgtatttgtcttatgggcacatggctttgagttgac
atctatgaagtattttgtgaaaataggacctgagcgcacctgttgtctatgtgatagacgtgccacatgcttttccactgcttcagacacttatg

cctgttggcatcattctattggatttgattacgtctataatccgtttatgattgatgttcaacaatggggttttacaggtaacctacaaagcaacc
atgatctgtattgtcaagtccatggtaatgcacatgtagctagttgtgatgcaatcatgactaggtgtctagctgtccacgagtgctttgttaag
cgtgttgactggactattgaatatcctataattggtgatgaactgaagattaatgcggcttgtagaaaggttcaacacatggttgttaaagctg
cattattagcagacaaattcccagttcttcacgacattggtaaccctaaagctattaagtgtgtacctcaagctgatgtagaatggaagttctat
gatgcacagccttgtagtgacaaagcttataaaatagaagaattattctattcttatgccacacattctgacaaattcacagatggtgtatgcc
tattttggaattgcaatgtcgatagatatcctgctaattccattgtttgtagatttgacactagagtgctatctaaccttaacttgcctggttgtga
tggtggcagtttgtatgtaaataaacatgcattccacacaccagcttttgataaaagtgcttttgttaatttaaaacaattaccatttttctattac
tctgacagtccatgtgagtctcatggaaaacaagtagtgtcagatatagattatgtaccactaaagtctgctacgtgtataacacgttgcaatt
taggtggtgctgtctgtagacatcatgctaatgagtacagattgtatctcgatgcttataacatgatgatctcagctggctttagcttgtgggttt
acaaacaatttgatacttataacctctggaacacttttacaagacttcagagtttagaaaatgtggcttttaatgttgtaaataagggacacttt
gatggacaacagggtgaagtaccagtttctatcattaataacactgtttacacaaaagttgatggtgttgatgtagaattgtttgaaaataaaa
caacattacctgttaatgtagcatttgagctttgggctaagcgcaacattaaaccagtaccagaggtgaaaatactcaataatttgggtgtgg
acattgctgctaatactgtgatctgggactacaaaagagatgctccagcacatatatctactattggtgtttgttctatgactgacatagccaag
aaaccaactgaaacgatttgtgcaccactcactgtcttttttgatggtagagttgatggtcaagtagacttatttagaaatgcccgtaatggtgt
tcttattacagaaggtagtgttaaaggtttacaaccatctgtaggtcccaaacaagctagtcttaatggagtcacattaattggagaagccgta
aaaacacagttcaattattataagaaagttgatggtgttgtccaacaattacctgaaacttactttactcagagtagaaatttacaagaattta
aacccaggagtcaaatggaaattgatttcttagaattagctatggatgaattcattgaacggtataaattagaaggctatgccttcgaacatat
cgtttatggagattttagtcatagtcagttaggtggtttacatctactgattggactagctaaacgttttaaggaatcaccttttgaattagaaga
ttttattcctatggacagtacagttaaaaactatttcataacagatgcgcaaacaggttcatctaagtgtgtgtgttctgttattgatttattactt
gatgattttgttgaaataataaaatcccaagatttatctgtagtttctaaggttgtcaaagtgactattgactatacagaaatttcatttatgcttt
ggtgtaaagatggccatgtagaaacattttacccaaaattacaatctagtcaagcgtggcaaccgggtgttgctatgcctaatctttacaaaa
tgcaaagaatgctattagaaaagtgtgaccttcaaaattatggtgatagtgcaacattacctaaaggcataatgatgaatgtcgcaaaatat
actcaactgtgtcaatatttaaacacattaacattagctgtaccctataatatgagagttatacattttggtgctggttctgataaaggagttgc
accaggtacagctgttttaagacagtggttgcctacgggtacgctgcttgtcgattcagatcttaatgactttgtctctgatgcagattcaacttt
gattggtgattgtgcaactgtacatacagctaataaatgggatctcattattagtgatatgtacgaccctaagactaaaaatgttacaaaaga
aaatgactctaaagagggtttttttcacttacatttgtgggtttatacaacaaaagctagctcttggaggttccgtggctataaagataacagaa
cattcttggaatgctgatctttataagctcatgggacacttcgcatggtggacagcctttgttactaatgtgaatgcgtcatcatctgaagcattt
ttaattggatgtaattatcttggcaaaccacgcgaacaaatagatggttatgtcatgcatgcaaattacatattttggaggaatacaaatccaa
ttcagttgtcttcctattctttatttgacatgagtaaatttccccttaaattaaggggtactgctgttatgtctttaaaagaaggtcaaatcaatga
tatgattttatctcttcttagtaaaggtagacttataattagagaaaacaacagagttgttatttctagtgatgttcttgttaacaactaaacgaa
caatgtttgtttttcttgttttattgccactagtctctagtcagtgtgttaatcttacaaccagaactcaattaccccctgcatacactaattctttca
cacgtggtgtttattaccctgacaaagttttcagatcctcagttttacattcaactcaggacttgttcttacctttcttttccaatgttacttggttcc
atgctatacatgtctctgggaccaatggtactaagaggtttgataaccctgtcctaccatttaatgatggtgtttattttgcttccactgagaagt
ctaacataataagaggctggatttttggtactactttagattcgaagacccagtccctacttattgttaataacgctactaatgttgttattaaag
tctgtgaatttcaattttgtaatgatccatttttgggtgtttattaccacaaaaacaacaaaagttggatggaaagtgagttcagagtttattcta
gtgcgaTGAGACG

pUC57-F6 Esp3I

CGTCTCAgcgaataattgcacttttgaatatgtctctcagccttttcttatggaccttgaaggaaaacagggtaatttcaaaaatcttaggg
aatttgtgtttaagaatattgatggttattttaaaatatattctaagcacacgcctattaatttagtgcgtgatctccctcagggtttttcggcttta
gaaccattggtagatttgccaataggtattaacatcactaggtttcaaactttacttgctttacatagaagttatttgactcctggtgattcttctt
caggttggacagctggtgctgcagcttattatgtgggttatcttcaacctaggacttttctattaaaatataatgaaaatggaaccattacagat
gctgtagactgtgcacttgaccctctctcagaaacaaagtgtacgttgaaatccttcactgtagaaaaaggaatctatcaaacttctaactta

gagtccaaccaacagaatctattgttagatttcctaatattacaaacttgtgcccttttggtgaagtttttaacgccaccagatttgcatctgttta
tgcttggaacaggaagagaatcagcaactgtgttgctgattattctgtcctatataattccgcatcattttccacttttaagtgttatggagtgtct
cctactaaattaaatgatctctgctttactaatgtctatgcagattcatttgtaattagaggtgatgaagtcagacaaatcgctccagggcaaa
ctggaaagattgctgattataattataaattaccagatgattttacaggctgcgttatagcttggaattctaacaatcttgattctaaggttggtg
gtaattataattacctgtatagattgtttaggaagtctaatctcaaaccttttgagagagatatttcaactgaaatctatcaggccggtagcaca
ccttgtaatggtgttgaaggttttaattgttactttcctttacaatcatatggtttccaacccactaatggtgttggttaccaaccatacagagtag
tagtactttcttttgaacttctacatgcaccagcaactgtttgtggacctaaaaagtctactaatttggttaaaaacaaatgtgtcaatttcaact
tcaatggtttaacaggcacaggtgttcttactgagtctaacaaaaagtttctgcctttccaacaatttggcagagacattgctgacactactgat
gctgtccgtgatccacagacacttgagattcttgacattacaccatgttcttttggtggtgtcagtgttataacaccaggaacaaatacttctaa
ccaggttgctgttctttatcaggatgttaactgcacagaagtccctgttgctattcatgcagatcaacttactcctacttggcgtgtttattctaca
ggttctaatgtttttcaaacacgtgcaggctgtttaataggggctgaacatgtcaacaactcatatgagtgtgacatacccattggtgcaggta
tatgcgctagttatcagactcagactaattctcctcggcgggcacgtagtgtagctagtcaatccatcattgcctacactatgtcacttggtgca
gaaaattcagttgcttactctaataactctattgccatacccacaaattttactattagtgttaccacagaaattctaccagtgtctatgaccaa
gacatcagtagattgtacaatgtacatttgtggtgattcaactgaatgcagcaatcttttgttgcaatatggcagtttttgtacacaattaaacc
gtgctttaactggaatagctgttgaacaagacaaaaacacccaagaagtttttgcacaagtcaaacaaatttacaaaacaccaccaattaaa
gattttggtggttttaatttttcacaaatattaccagatccatcaaaaccaagcaagaggtcatttattgaagatctacttttcaacaaagtgac
acttgcagatgctggcttcatcaaacaatatggtgattgccttggtgatattgctgctagagacctcatttgtgcacaaaagtttaacggcctta
ctgttttgccacctttgctcacagatgaaatgattgctcaatacacttctgcactgttagcgggtacaatcacttctggttggacctttggtgcag
gtgctgcattacaaataccatttgctatgcaaatggcttataggtttaatggtattggagttacacagaatgttctctatgagaaccaaaaattg
attgccaaccaatttaatagtgctattggcaaaattcaagactcactttcttccacagcaagtgcacttggaaaacttcaagatgtggtcaacc
aaaatgcacaagctttaaacacgcttgttaaacaacttagctccaattttggtgcaatttcaagtgttttaaatgatatcctttcacgtcttgaca
aagttgaggctgaagtgcaaattgataggttgatcacaggcagacttcaaagtttgcagacatatgtgactcaacaattaattagagctgcag
aaatcagagcttctgctaatcttgctgctactaaaatgtcagagtgtgtacttggacaatcaaaaagagttgatttttgtggaaagggctatca
tcttatgtccttccctcagtcagcacctcatggtgtagtcttcttgcatgtgacttatgtccctgcacaagaaaagaacttcacaactgctcctgc
catttgtcatgatggaaaagcacactttcctcgtgaaggtgtctttgtttcaaatggcacacactggtttgtaacacaaaggaatttttatgaac
cacaaatcattactacagacaacacatttgtgtctggtaactgtgatgttgtaataggaattgtcaacaacacagtttatgatcctttgcaacct
gaattagactcattcaaggaggagttagataaatattttaagaatcatacatcaccagatgttgatttaggtgacatctctggcattaatgcttc
agttgtaaacattcaaaaagaaattgaccgcctcaatgaggttgccaagaatttaaatgaatctctcatcgatctccaagaacttggaaagta
tgagcagtatataaaatggccatggtacatttggctaggttttatagctggcttgattgccatagtaatggtgacaattatgctttgctgtatga
ccagttgctgtagttgtctcaagggctgttgttcttgtggatcctgctgcaaatttgatgaagacgactctgagccagtgctcaaaggagtcaa
attacattacacataaacgaacttatggatttgtttatgagaatcttcacaattggaactgtaactttgaagcaaggtgaaatcaaggatgcta
ctccttcagattttgttcgcgctactgcaacgataccgatacaagcctcactccctttcggatggcttattgttggcgttgcacttcttgctgttttt
cagagcgcttccaaaatcataaccctcaaaaagagatggcaactagcactctccaagggtgttcactttgtttgcaacttgctgttgtttgt
aacagtttactcacaccttttgctcgttgctgctggccttgaagccccttttctctatctttatgctttagtctacttcttgcagagtataaactttgt
aagaataataatgaggctttggctttgctggaaatgccgttccaaaaacccattactttatgatgccaactattttctttgctggcatactaattg
ttacgactattgtataccttacaatagtgtaacttcttcaattgtcattacttcaggtgatggcacaacaagtcctatttctgaacatgactacca
gattggtggttatactgaaaaatgggaatctggagtaaaagactgtgttgtattacacagttacttcacttcagactattaccagctgtactca
actcaattgagtacagacactggtgttgaacatgttaccttcttcatctacaataaaattgttgatgagcctgaagaacatgtccaaattcaca
caatcgacggttcatccggagttgttaatccagtaatggaaccaatttatgatgaaccgacgacgactactagcgtgcctttgtaagcacaag
ctgatgagtacgaacttatgtactcattcgtttcggaagagacaggtacgttaatagttaatagcgtacttctttttcttgctttcgtggtattcttg
ctagttacactTGAGACG

pCC1-F7 Esp3I

CGTCTCAcactagccatccttactgcgcttcgattgtgtgcgtactgctgcaatattgttaacgtgagtcttgtaaaaccttctttttacg
tttactctcgtgttaaaaatctgaattcttctagagttcctgatcttctggtctaaacgaactaaatattatattagtttttctgtttggaact
ttaattttagccatggcagattccaacggtactattaccgttgaagagcttaaaaagctccttgaacaatggaacctagtaataggtttc
ctattccttacatggatttgtcttctacaatttgcctatgccaacaggaataggtttttgtatataattaagttaattttcctctggctgttat
ggccagtaactttagcttgttttgtgcttgctgctgtttacagaataaattggatcaccggtggaattgctatcgcaatggcttgtcttgta
ggcttgatgtggctcagctacttcattgcttctttcagactgtttgcgcgtacgcgttccatgtggtcattcaatccagaaactaacattctt
ctcaacgtgccactccatggcactattctgaccagaccgcttctagaaagtgaactcgtaatcggagctgtgatccttcgtggacatctt
cgtattgctggacaccatctaggacgctgtgacatcaaggacctgcctaaagaaatcactgttgctacatcacgaacgctttcttattac
aaattgggagcttcgcagcgtgtagcaggtgactcaggttttgctgcatacagtcgctacaggattggcaactataaattaaacacag
accattccagtagcagtgacaatattgctttgcttgtacagtaagtgacaacagatgtttcatctcgttgactttcaggttactatagcag
agatattactaattattatgaggacttttaaagtttccatttggaatcttgattacatcataaacctcataattaaaaatttatctaagtca
ctaactgagaataaatattctcaattagatgaagagcaaccaatggagattgattaaacgaacatgaaaattattcttttcttggcact
gataacactcgctacttgtgagctttatcactaccaagagtgtgttagaggtacaacagtacttttaaaagaaccttgctcttctggaac
atacgagggcaattcaccatttcatcctctagctgataacaaatttgcactgacttgctttagcactcaatttgcttttgcttgtcctgacg
gcgtaaaacacgtctatcagttacgtgccagatcagtttcacctaaactgttcatcagacaagaggaagttcaagaactttactctcca
atttttcttattgttgcggcaatagtgtttataacactttgcttcacactcaaaagaaagacagaatgattgaactttcattaattgacttct
atttgtgctttttagcctttctgctattccttgttttaattatgcttattatcttttggttctcacttgaactgcaagatcataatgaaacttgtc
acgcctaaacgaacatgaaatttcttgttttcttaggaatcatcacaactgtagctgcatttcaccaagaatgtagtttacagtcatgtac
tcaacatcaaccatatgtagttgatgacccgtgtcctattcacttctattctaaatggtatattagagtaggagctagaaaatcagcacc
tttaattgaattgtgcgtggatgaggctggttctaaatcacccattcagtacatcgatatcggtaattatacagtttcctgtttaccttttac
aattaattgccaggaacctaaattgggtagtcttgtagtgcgttgttcgttctatgaagactttttagagtatcatgacgttcgtgttgtttt
agatttcatctaaacgaacaaactaaaatgtctgataatggaccccaaaatcagcgaaatgcaccccgcattacgtttggtggaccct
cagattcaactggcagtaaccagaatggagaacgcagtggggcgcgatcaaaacaacgtcggccccaaggtttacccaataatact
gcgtcttggttcaccgctctcactcaacatggcaaggaagaccttaaattccctcgaggacaaggcgttccaattaacaccaatagca
gtccagatgaccaaattggctactaccgaagagctaccagacgaattcgtggtggtgacggtaaaatgaaagatctcagtccaagat
ggtatttctactacctaggaactgggccagaagctggacttccctatggtgctaacaaagacggcatcatatgggttgcaactgaggg
agccttgaatacaccaaaagatcacattggcacccgcaatcctgctaacaatgctgcaatcgtgctacaacttcctcaaggaacaaca
ttgccaaaaggcttctacgcagaagggagcagaggcggcagtcaagcctcttctcgttcctcatcacgtagtcgcaacagttcaagaa
attcaactccaggcagcagtaggggaacttctcctgctagaatggctggcaatggcggtgatgctgctcttgctttgctgctgcttgaca
gattgaaccagcttgagagcaaaatgtctggtaaaggccaacaacaacaaggccaaactgtcactaagaaatctgctgctgaggctt
ctaagaagcctcggcaaaaacgtactgccactaaagcatacaatgtaacacaagctttcggcagacgtggtccagaacaaacccaa
ggaaattttggggaccaggaactaatcagacaaggaactgattacaaacattggccgcaaattgcacaatttgcccccagcgcttca
gcgttcttcggaatgtcgcgcattggcatggaagtcacaccttcgggaacgtggttgacctacacaggtgccatcaaattggatgaca
aagatccaaatttcaaagatcaagtcattttgctgaataagcatattgacgcatacaaaacattcccaccaacagagcctaaaaagga
caaaaagaagaaggctgatgaaactcaagccttaccgcagagacagaagaaacagcaaactgtgactcttcttcctgctgcagattt
ggatgatttctccaaacaattgcaacaatccatgagcagtgctgactcaactcaggcctaaactcatgcagaccacacaaggcagat
gggctatataaacgttttcgcttttccgtttacgatatatagtctactcttgtgcagaatgaattctcgtaactacatagcacaagtagatg
tagttaactttaatctcacatagcaatctttaatcagtgtgtaacattagggaggacttgaaagagccaccacattttcaccgaggccac
gcggagtacgatcgagtgtacagtgaacaatgctagggagagctgcctatatggaagagccctaatgtgtaaaattaattttagtagt
gctatccccatgtgattttaatagcttcttaggagaatgacaaaaaaaaaaaaaaaaaaaaaaaaaaaaaaaaaTGAGACG
