## Supplementary Figure S1-S15,S17 for "Nucleocapsid mutation R203K/G204R increases the infectivity, fitness and virulence of SARS-CoV-2"

A

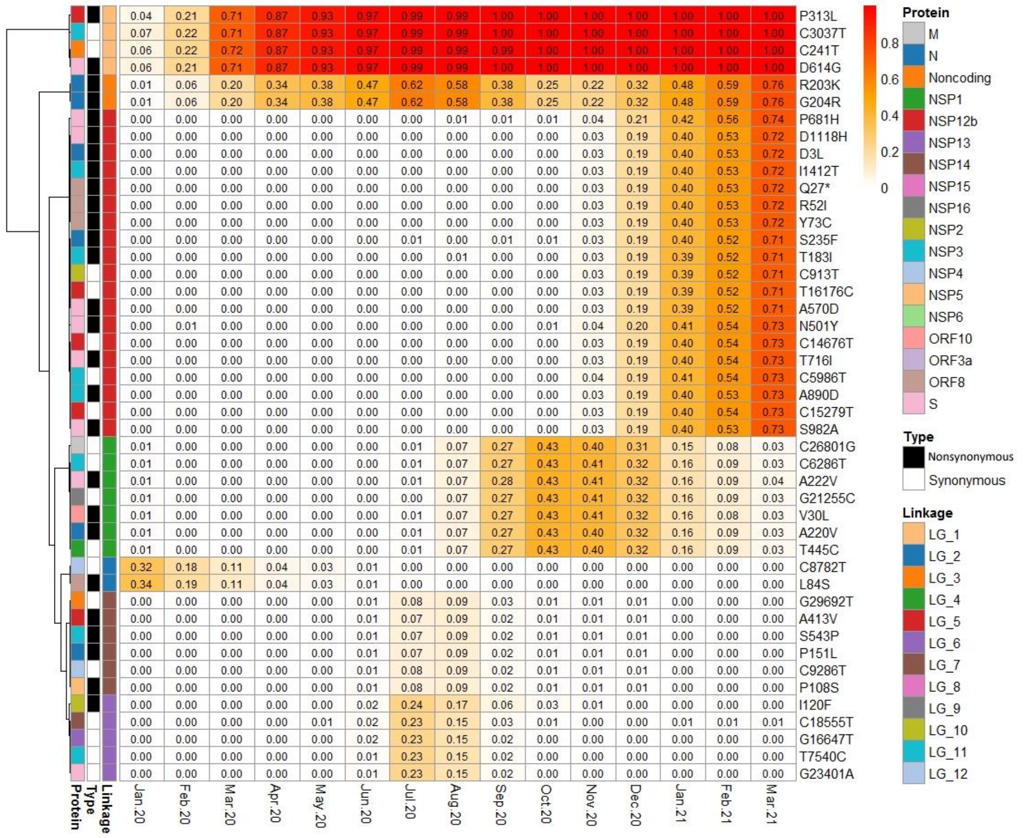
P313L
C3037T
C241T
D614G
R203K
G204R
P681H
D1118H
D3L
I1412T
Q27*
R52I
Y73C
S235F
T183I
C913T
T16176C
A570D
N501Y
C14676T
T716I
C5986T
A890D
C15279T
S982A
C26801G
C6286T
A222V
G21255C
V30L
A220V
T445C
C8782T
L84S
G29692T
A413V
S543P
P151L
C9286T
P108S
I120F
C18555T
G16647T
T7540C
G23401A
Protein
Type
Linkage
Jan.20
Feb.20
Mar.20
Apr.20
May.20
Jun.20
Jul.20
Aug.20
Sep.20
Oct.20
Nov.20
Dec.20
Jan.21
Feb.21
Mar.21
0.8
0.6
0.4
0.2
0
Protein
M
N
Noncoding
NSP1
NSP12b
NSP13
NSP14
NSP15
NSP16
NSP2
NSP3
NSP4
NSP5
NSP6
ORF10
ORF3a
ORF8
S
Type
Nonsynonymous
Synonymous
Linkage
LG_1
LG_2
LG_3
LG_4
LG_5
LG_6
LG_7
LG_8
LG_9
LG_10
LG_11
LG_12

B

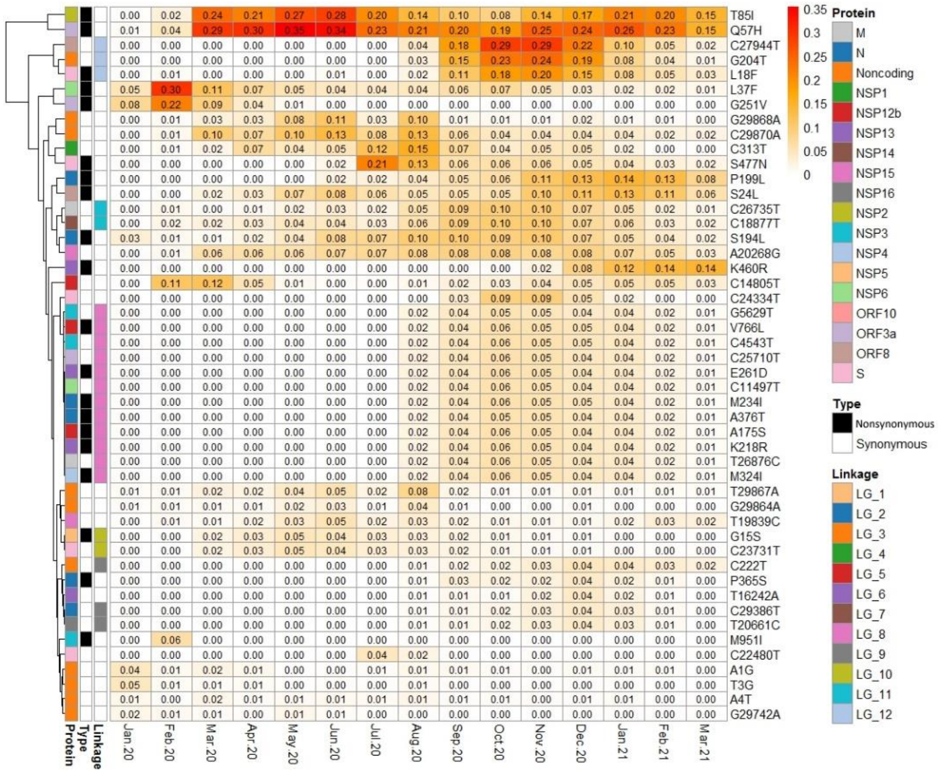
T85I
Q57H
C27944T
G204T
L18F
L37F
G251V
G29868A
C29870A
C313T
S477N
P199L
S24L
C26735T
C18877T
S194L
A20268G
K460R
C14805T
C24334T
G5629T
V766L
C4543T
C25710T
E261D
C11497T
M234I
A376T
A175S
K218R
T26876C
M324I
T29867A
G29864A
T19839C
G15S
C23731T
C222T
P365S
T16242A
C29386T
T20661C
M951I
C22480T
A1G
T3G
A4T
G29742A
Protein
Type
Linkage
Jan.20
Feb.20
Mar.20
Apr.20
May.20
Jun.20
Jul.20
Aug.20
Sep.20
Oct.20
Nov.20
Dec.20
Jan.21
Feb.21
Mar.21
0.35
0.3
0.25
0.2
0.15
0.1
0.05
0
Protein
M
N
Noncoding
NSP1
NSP12b
NSP13
NSP14
NSP15
NSP16
NSP2
NSP3
NSP4
NSP5
NSP6
ORF10
ORF3a
ORF8
S
Type
Nonsynonymous
Synonymous
Linkage
LG_1
LG_2
LG_3
LG_4
LG_5
LG_6
LG_7
LG_8
LG_9
LG_10
LG_11
LG_12

Figure S1. The IF changes of 96 mutations from Jan-2020 to Mar-2021. A refers to mutations in LG_1 to LG7. B refers to mutations in LG_8 to LG_12 and other singleton mutations. The depth of color indicates the value of IF. There are annotation rows showing the affected protein and the mutation type (synonymous and nonsynonymous).

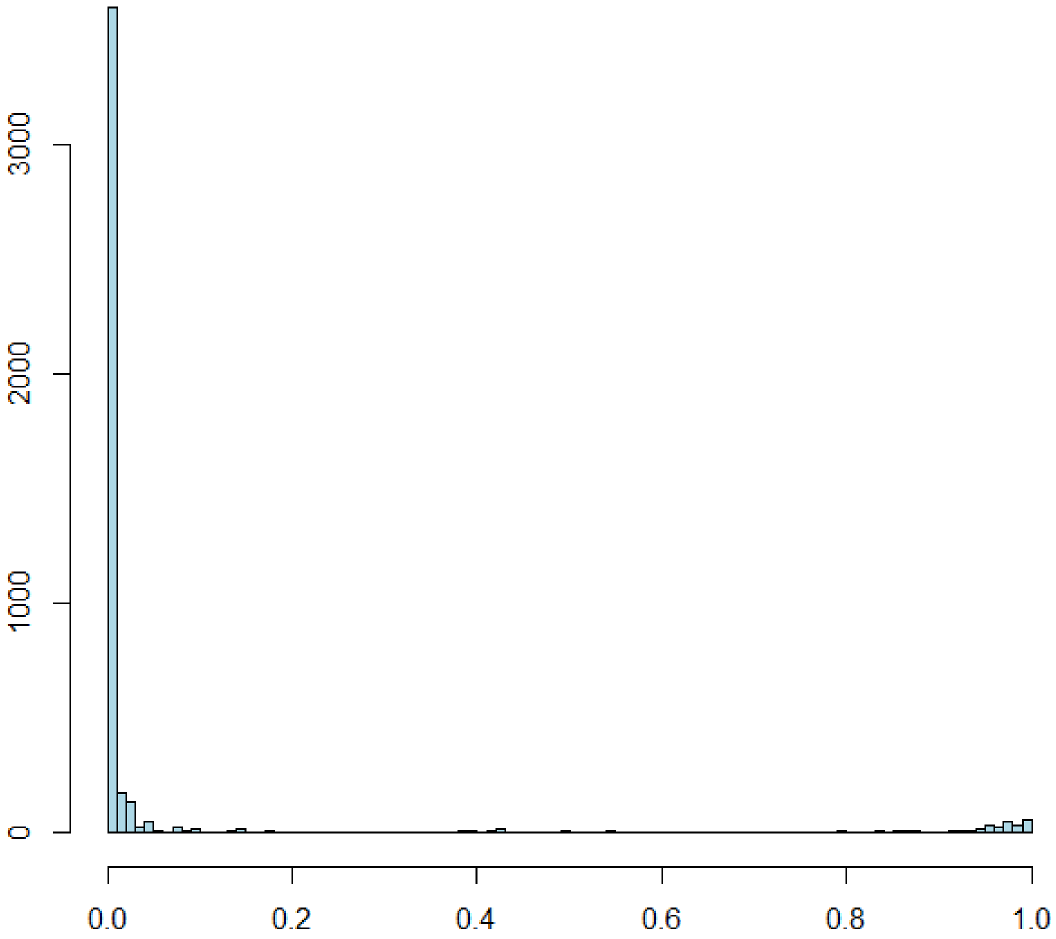


Figure S2. The distribution of $\rho^2$ for pairs of mutations shown in Figure S1.

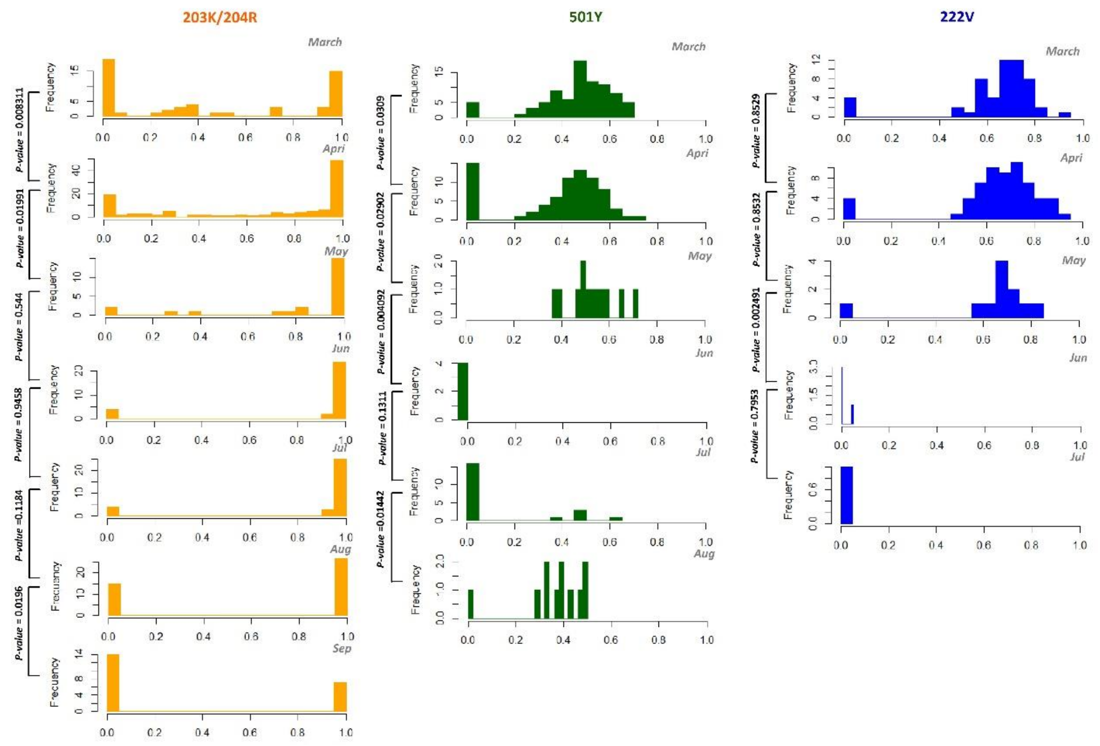


Figure S3. Changes in the distribution of IFs along time for 203K/204R, 501Y and 222V. The iSNV analyses are based on the SARS-CoV-2 raw-sequence data collected in the United Kingdom. X-axis is the IF, whose value ranges from 0 to 1 (fixed), and the Y-axis refers to the counts of samples with different IF values. Results of Wilcoxon test for the difference between two distributions are shown at the left of each figure.

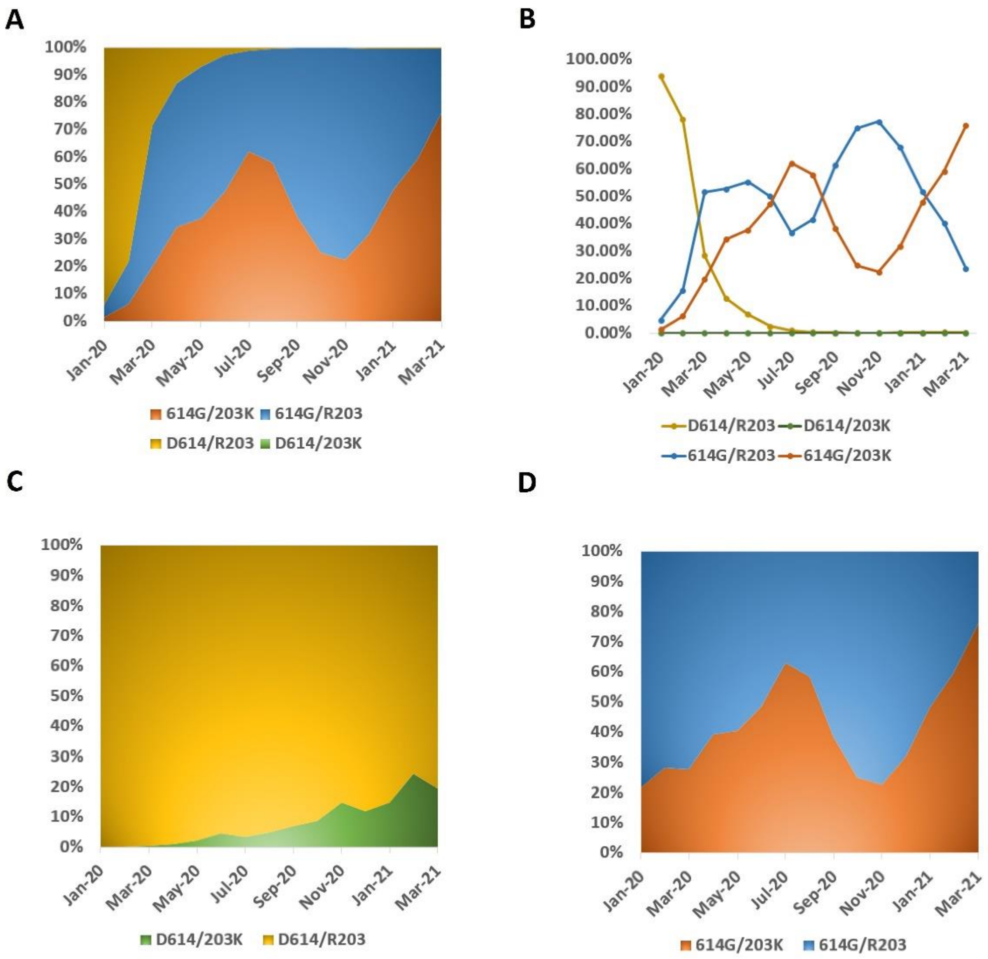


Figure S4. The IF changes of four combinations, D614/R203, D614/203K, 614G/R203 and 614G/203K. A, C and D are the percentage accumulated area map of the four combinations, D614/R203 vs D614/203K, and 614G/R203 vs 614G/203K. B is the line chart according to A.

**A**

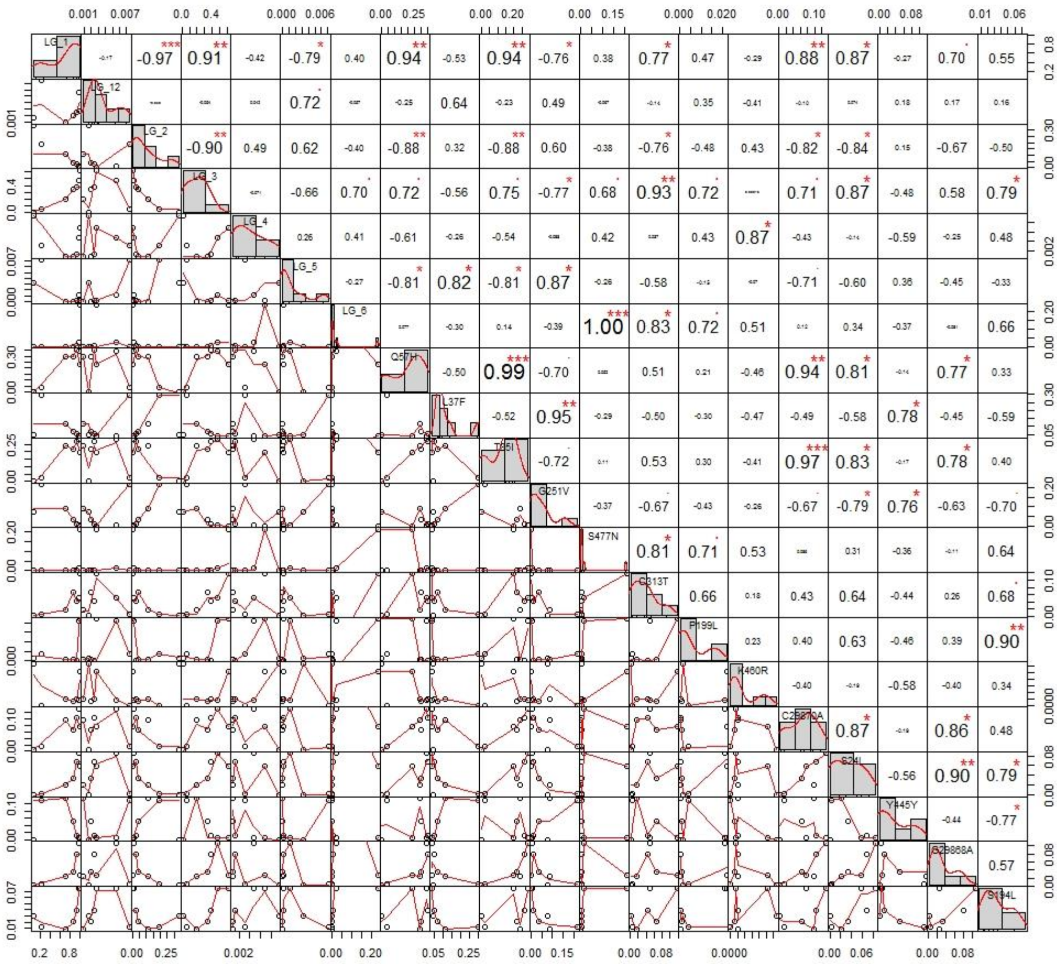

LG_1
LG_12
LG_2
LG_3
LG_4
LG_5
LG_6
Q57H
L37F
T85I
G251V
S477N
C313T
P199L
K460R
S24L
Y445Y
S194L

B

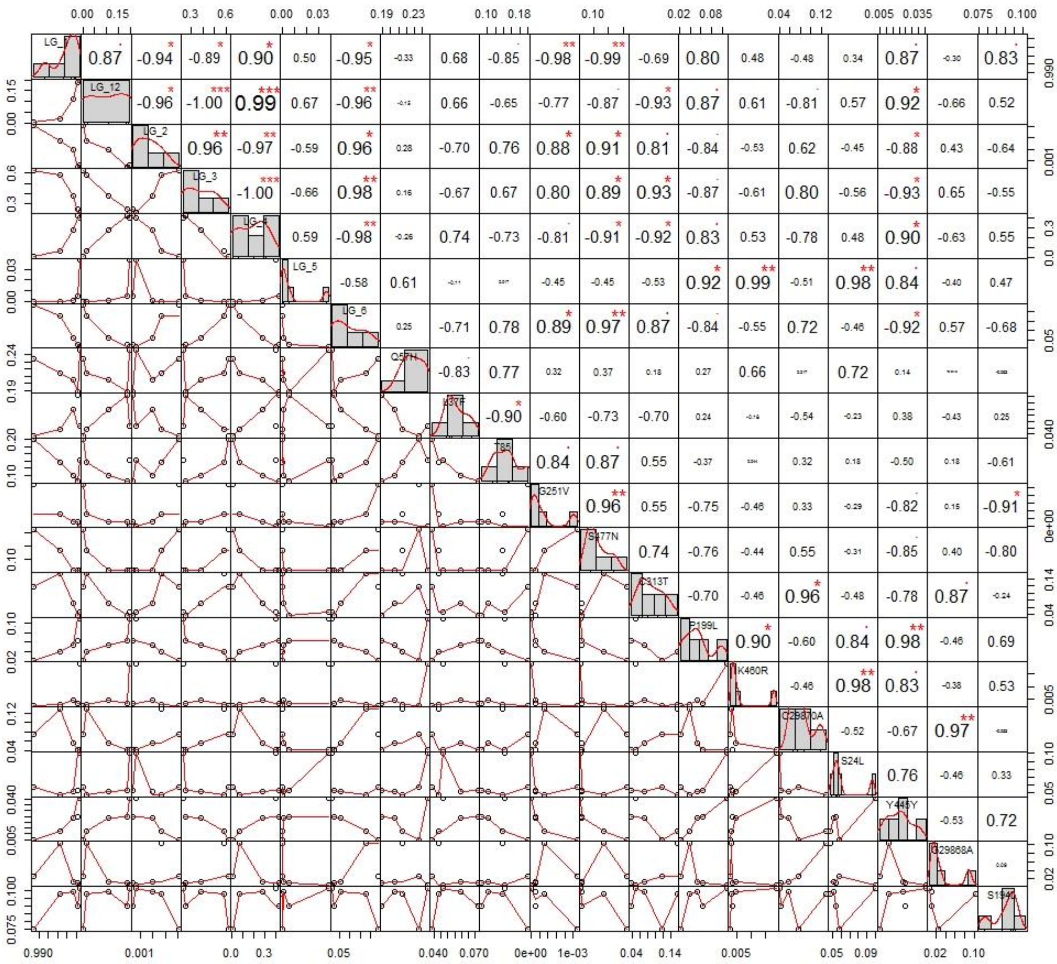

**C**

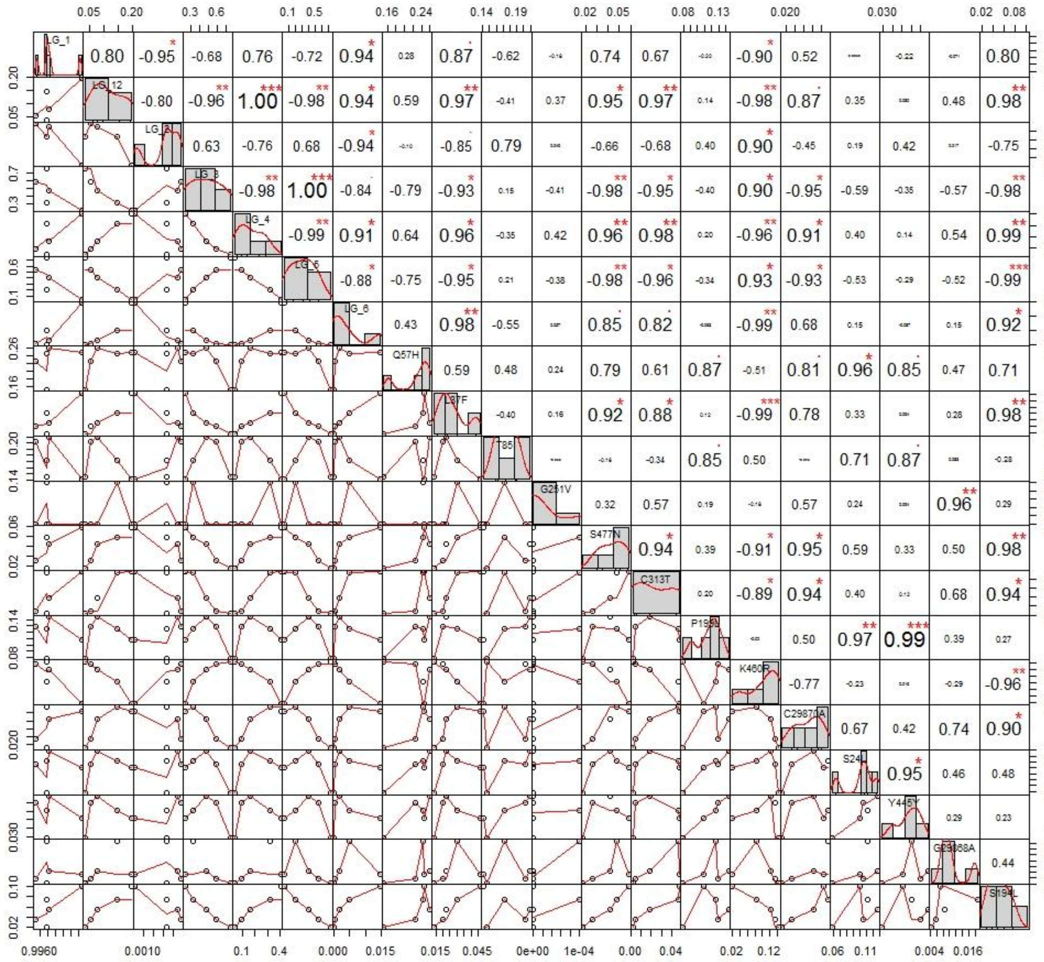

Figure S5. Correlation tests between pairs of LGs/mutations. Tests are performed in three time intervals, I1 (A), D (B) and I2 (C), as shown in Figure 1B. “*”, “**” and “***” denote P-value < 0.1, P-value < 0.05 and P-value < 0.01, respectively.

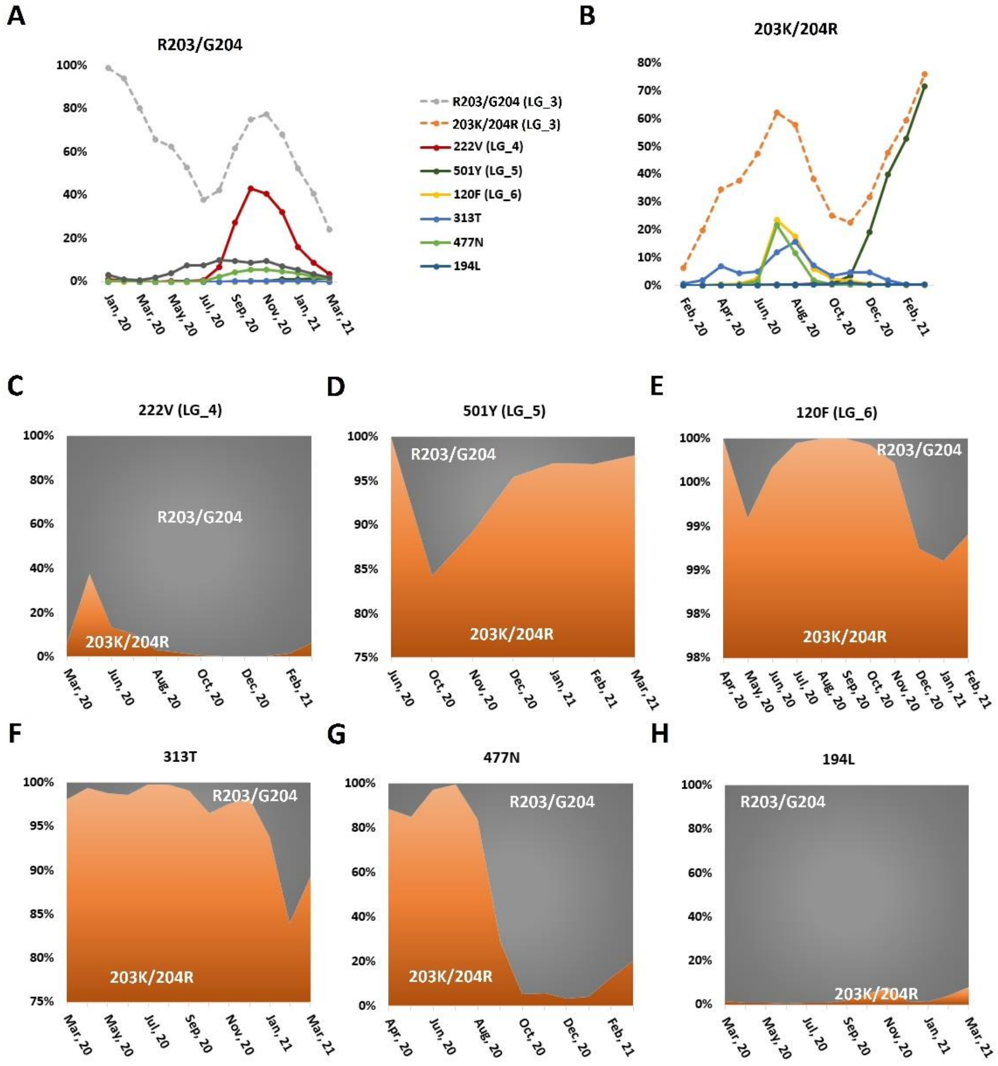


Figure S6. Comparison of IF tracks for the six mutants with significant correlation with LG_3. A and B are the IFs of the six mutants (real lines) in R203/G204 and 203K/204R variants, respectively. In A and B, dotted lines denote the IF changes of R203/G204 and 203K/204R variants, respectively. C to H are the ratios of R203/G204 and 203K/204R in months in the six mutants.

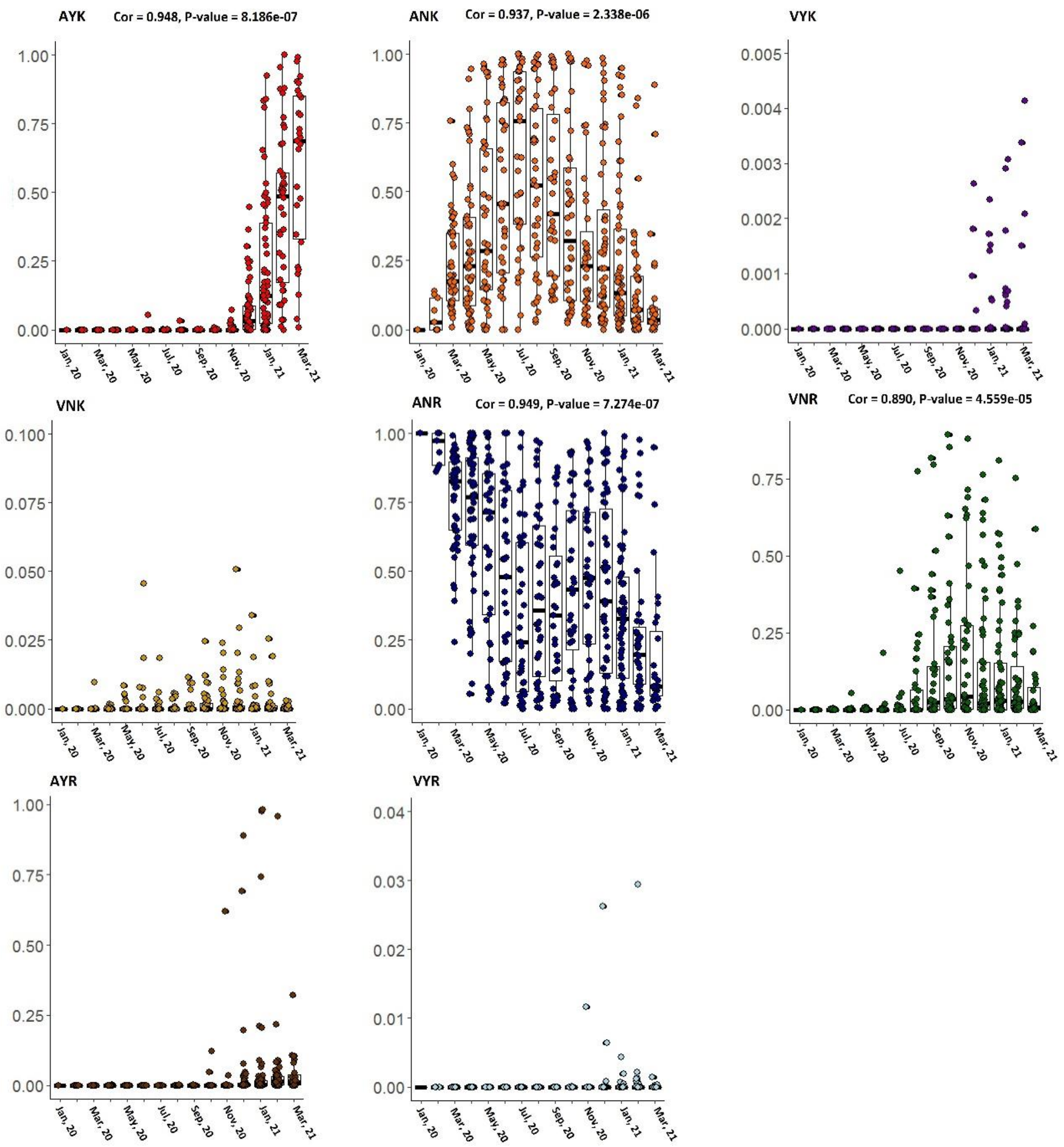


Figure S7. IFs in countries for the 8 lineages shown in Figure 2. For the four dominant lineages shown in Figure 2A, we calculated the correlation between the median IFs in countries and the IFs in the world shown in Figure 2A. The results display at the top.

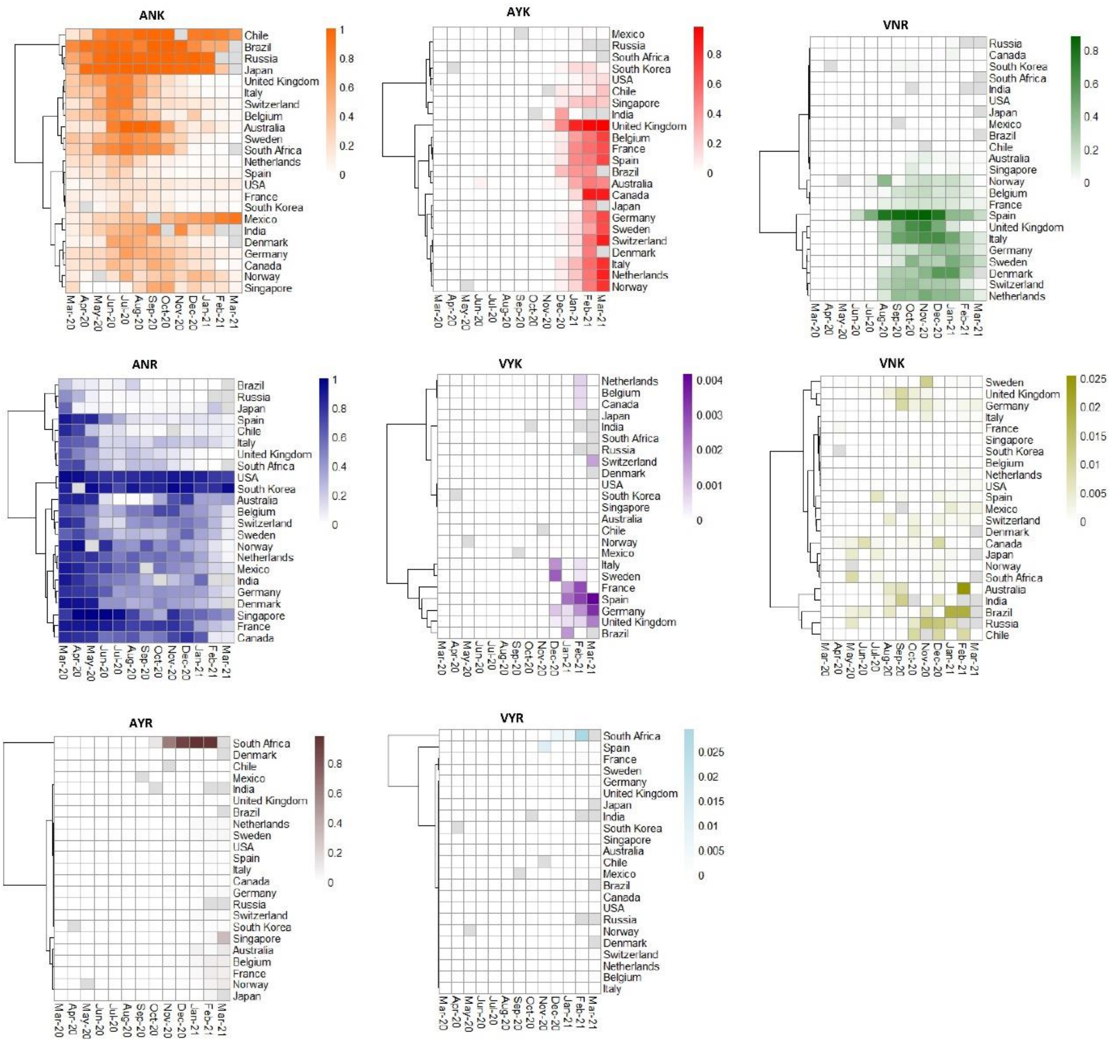


Figure S8. Heatmaps showing the IF changes in countries for the eight lineages shown in Figure 2. We did not showed the countries without an identification IF > 0 in any month, mostly for the insufficiency of samples. The empty IFs are colored grey.

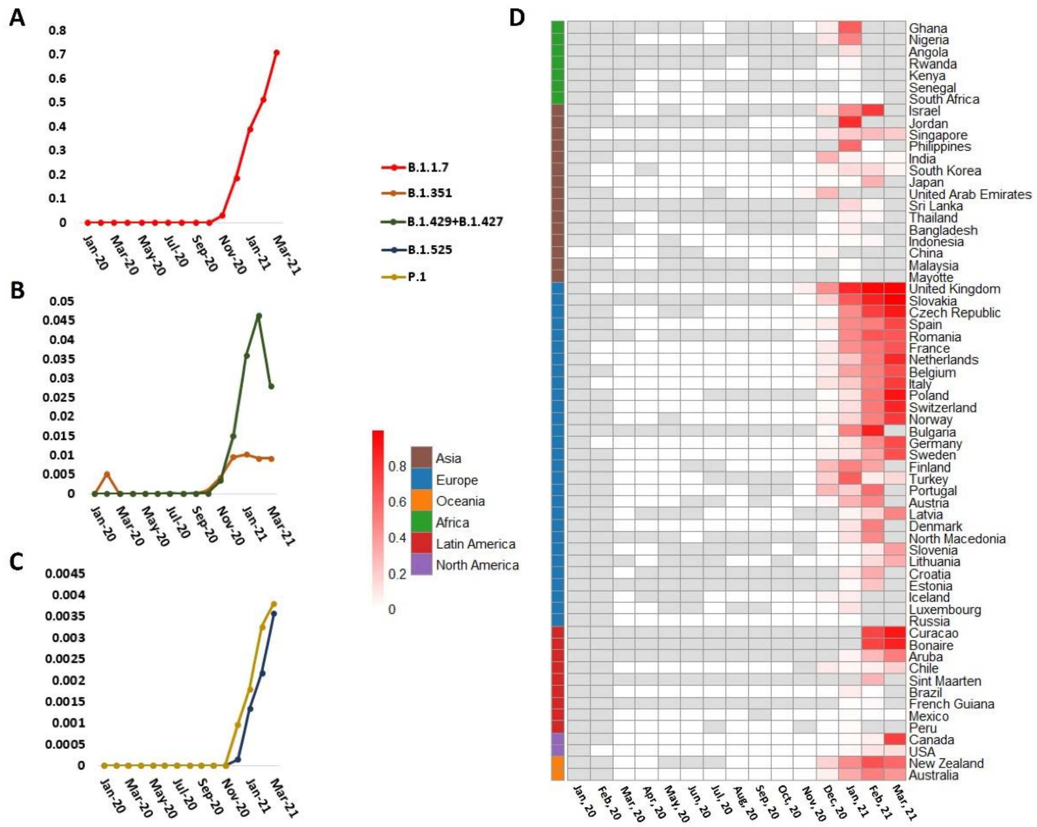


Figure S9. The IF change of fast spread lineages. A to C is the IF change of five lineages in the world. D is the change of IF in countries for the lineage B.1.1.7. In D, continents are marked in the annotation row at the left.

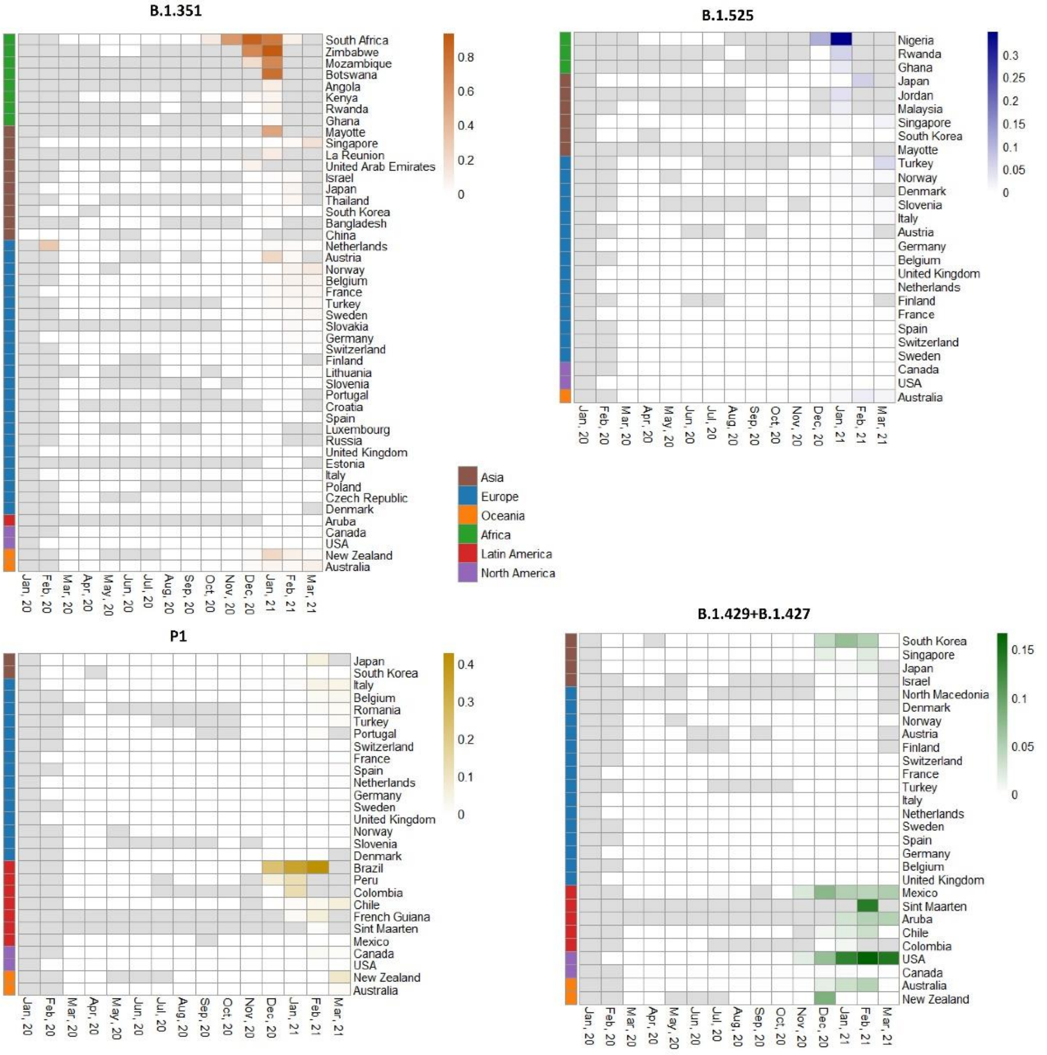


Figure S10. The IF change in countries for four lineages. Other legends follow Figure S9.

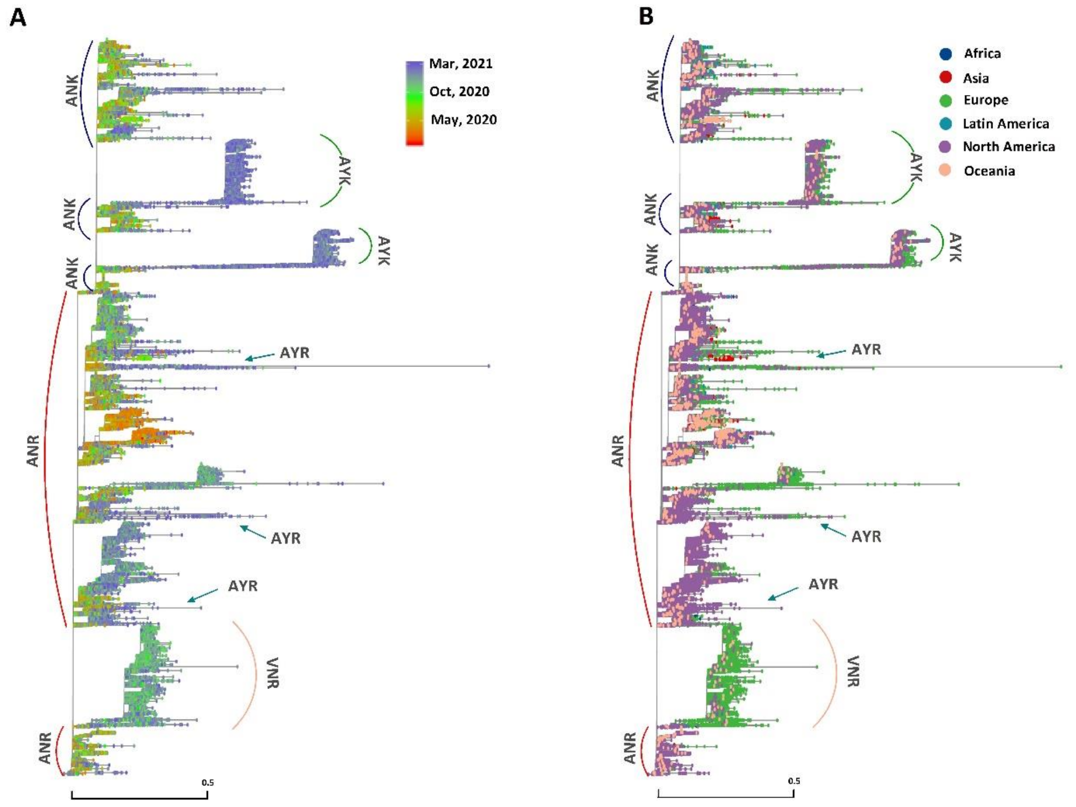


Figure S11. Distribution of the collection months and continents in the SARS-CoV-2 phylogenetic tree. In A, the colors of tip nodes denote the collection months. In B, the colors of tip nodes denote the collection continents. Lineages are marked.

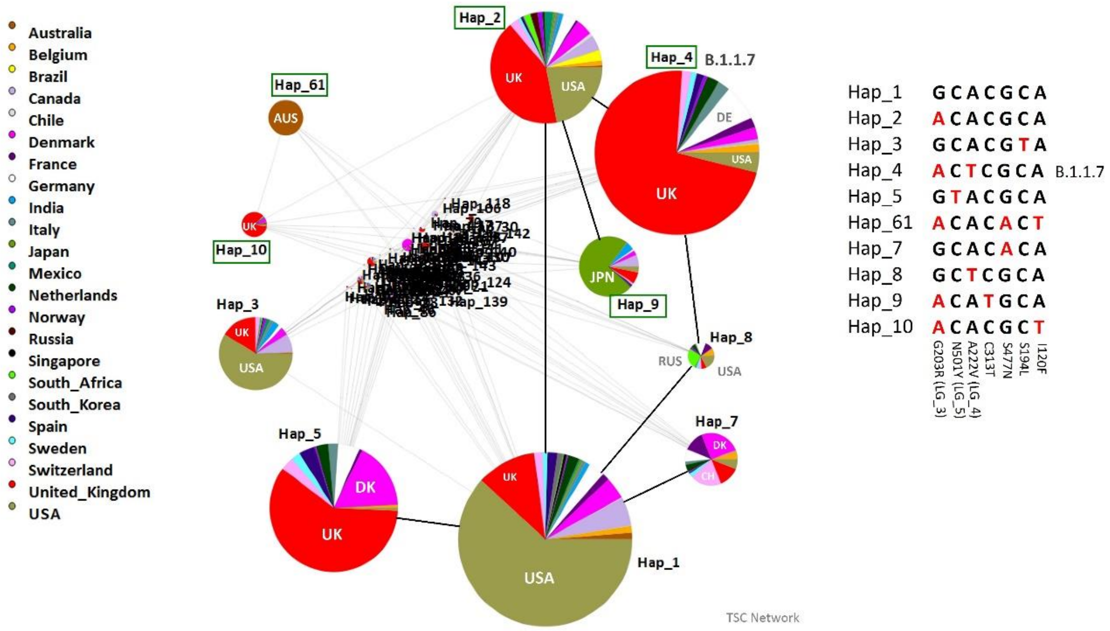


Figure S12. A Network of haplotypes constructed by 7 LGs/mutations. The geological positions are differentiated by different colors. The sequences of the haplotypes are shown in the right, with mutants marked in red.

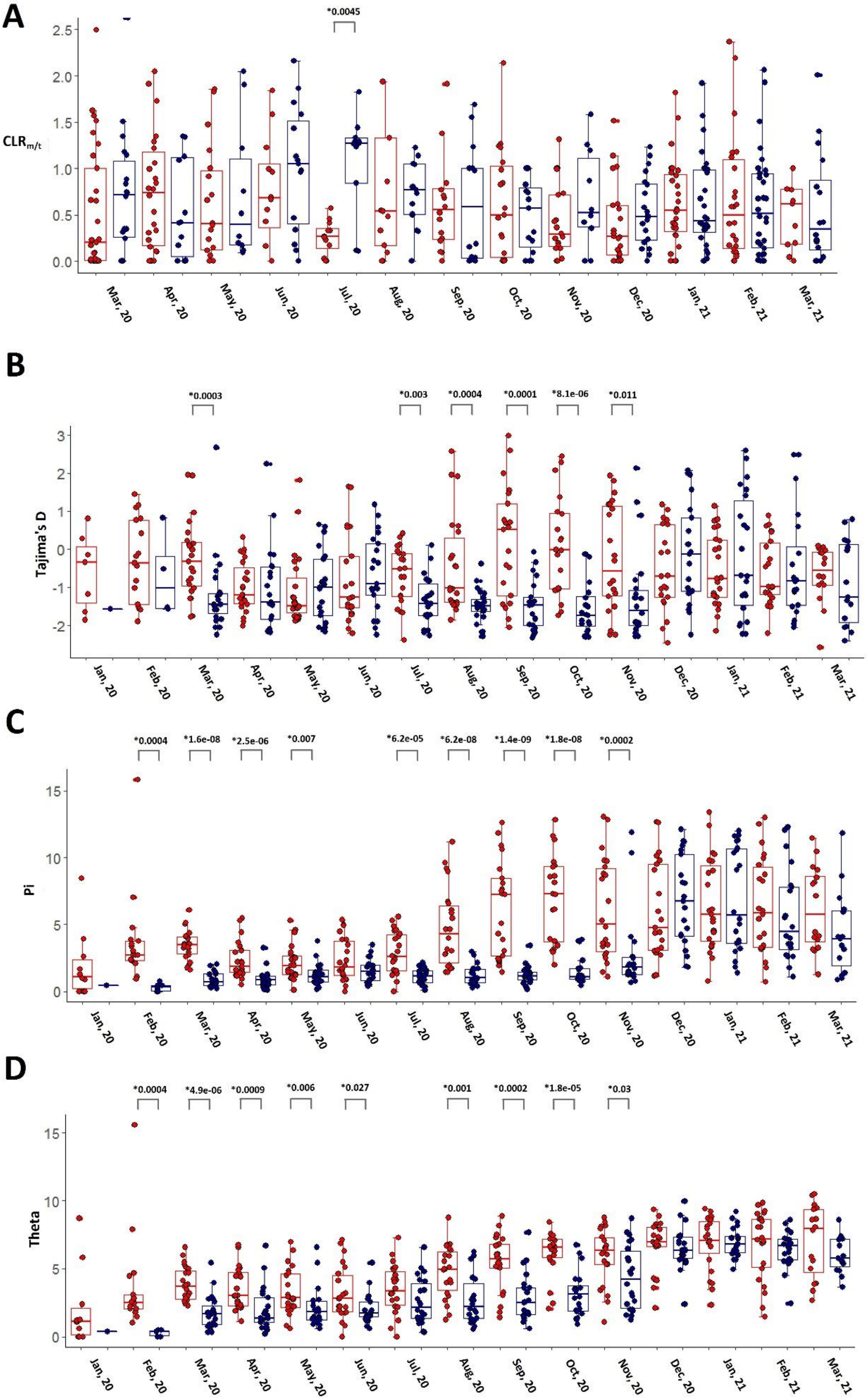
A
*0.0045
CLR$_{m/t}$
2.5
2.0
1.5
1.0
0.5
0.0
Mar, 20
Apr, 20
May, 20
Jun, 20
Jul, 20
Aug, 20
Sep, 20
Oct, 20
Nov, 20
Dec, 20
Jan, 21
Feb, 21
Mar, 21
B
*0.0003
*0.003
*0.0004
*0.0001
*8.1e-06
*0.011
Tajima's D
3
2
1
0
-1
-2
Jan, 20
Feb, 20
Mar, 20
Apr, 20
May, 20
Jun, 20
Jul, 20
Aug, 20
Sep, 20
Oct, 20
Nov, 20
Dec, 20
Jan, 21
Feb, 21
Mar, 21
C
*0.0004
*1.6e-08
*2.5e-06
*0.007
*6.2e-05
*6.2e-08
*1.4e-09
*1.8e-08
*0.0002
Pi
15
10
5
0
Jan, 20
Feb, 20
Mar, 20
Apr, 20
May, 20
Jun, 20
Jul, 20
Aug, 20
Sep, 20
Oct, 20
Nov, 20
Dec, 20
Jan, 21
Feb, 21
Mar, 21
D
*0.0004
*4.9e-06
*0.0009
*0.006
*0.027
*0.001
*0.0002
*1.8e-05
*0.03
Theta
15
10
5
0
Jan, 20
Feb, 20
Mar, 20
Apr, 20
May, 20
Jun, 20
Jul, 20
Aug, 20
Sep, 20
Oct, 20
Nov, 20
Dec, 20
Jan, 21
Feb, 21
Mar, 21

Figure S13. Selection signatures for R203K/G204R.

(A) Comparison of CLR $_{m/t}$ in months between R203/G204 (red) and 203K/204R variants (darkblue).

(B-D) Comparisons of Tajima's D (B), Pi (C) and Theta (D) in the whole genome of R203/G204 (colored in red) and 203K/204R strains (colored in darkblue). Comparisons with statistical significance are marked.

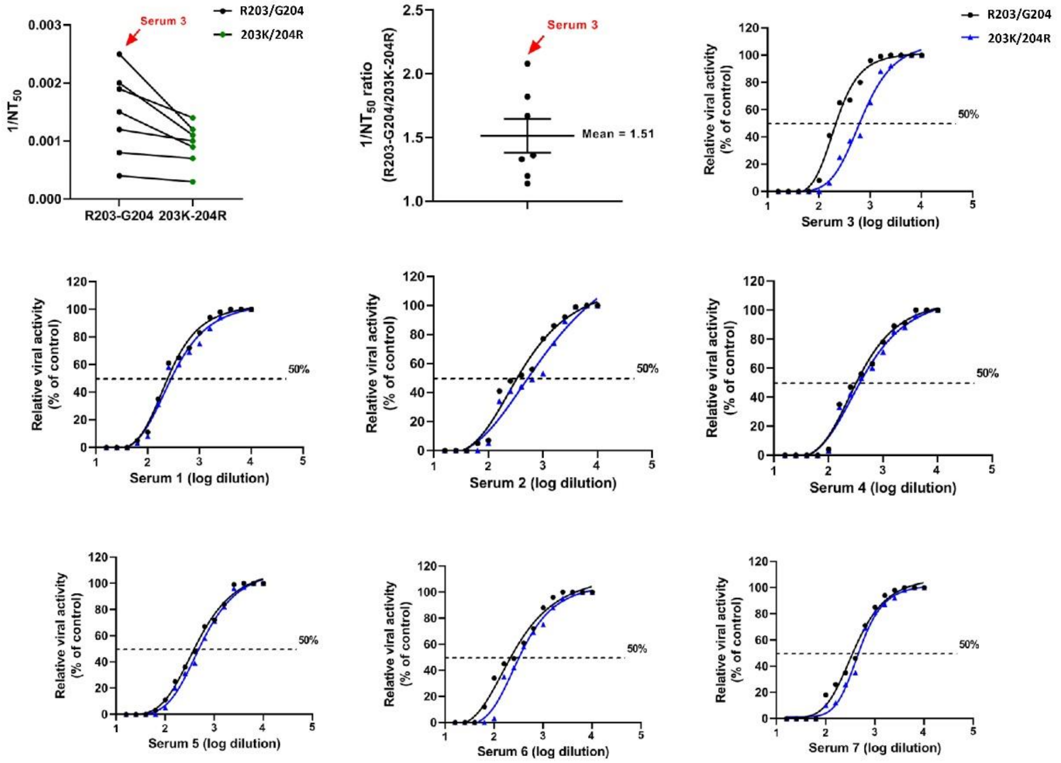


Figure S14. Stability of R203/G204 and 203K/204R viruses

(A) Neutralizing activities of hamster sera against R203/G204 and 203K/204R viruses with mNeonGreen reporter. The $1/NT_{50}$ values were plotted. Symbols represented sera from individual hamsters. (B) Ratio of $1/NT_{50}$ between R203/G204 and 203K/204R viruses. Symbols represented sera from individual hamsters. (C-I) Neutralizing curves of serum from individual hamsters. The solid line represented the fitted curve, and the dotted line indicated 50% viral inhibition. Data in (B) were represented as mean±s.e.m..

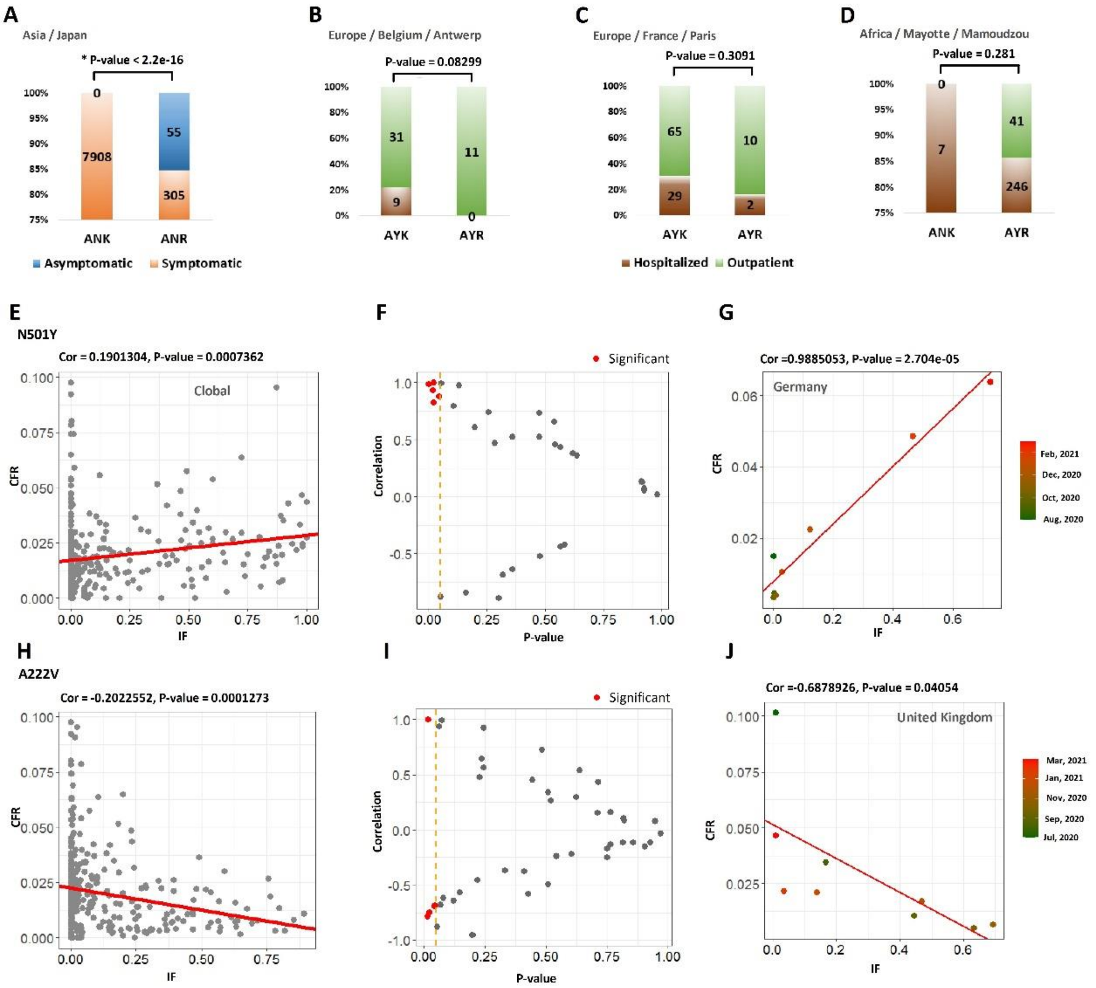


Figure S15. (A-D) Prediction on the clinical outcomes of ANK, ANR, AYK and AYR based on the clinical data collected in a small scale. (E-J) Correlation analysis results between the mutant IFs and CFR, for N501Y and A222V. Legends follow Figure 6.

Figure S16. The sequences of F1~F7 fragments and the restriction enzymes used for digestion and ligation. The Figure is too large and is put in an additional file.

F: 5’-GGCAGCAGTAAACGAACTTCTCCTGCTAGAATGGCTGGCAATGGCGGTGATGCTG-3’,

R: 5’-CACCGCCATTGCCAGCCATTCTAGCAGGAGAAGTTTTGCTACTGCTGCCTGGAGTTG-3

Figure S17. The primers used for overlap-extension PCR in the generation of R203K/G204R mutant virus. Nucleotides underlined are nucleotide substitutions. Nucleotides colored denote the reading frame (R203K/G204R mutation).
